# A phenotypic rescue approach identifies lineage regionalization defects in a mouse model of DiGeorge syndrome

**DOI:** 10.1101/2021.12.07.471208

**Authors:** Gabriella Lania, Monica Franzese, Adachi Noritaka, Marchesa Bilio, Annalaura Russo, Erika D’Agostino, Claudia Angelini, Robert G. Kelly, Antonio Baldini

## Abstract

*TBX1* is a key regulator of pharyngeal apparatus (PhAp) development. Vitamin B12 treatment partially rescues aortic arch patterning defects of *Tbx1^+/-^* embryos. Here we show that it also improves cardiac outflow tract septation and branchiomeric muscle anomalies of *Tbx1* hypomorphic mutants. At molecular level, the in vivo vB12 treatment let us to identify genes that were dysregulated by *Tbx1* haploinsufficiency and rescued by treatment. We found that SLUG, encoded by the rescued gene *Snai2*, identified a population of mesodermal cells that was partially overlapping with but distinct from ISL1+ and TBX1+ populations. In addition, SLUG+ cells were mislocalized and had a greater tendency to aggregate in *Tbx1^+/-^ and Tbx1^-/-^* embryos and vB12 treatment restore cellular distribution. Adjacent neural crest-derived mesenchymal cells, which do not express TBX1, were also affected, showing enhanced segregation from cardiopharyngeal mesodermal cells. We propose that TBX1 regulates cell distribution in core mesoderm and the arrangement of multiple lineages within the PhAp.

## INTRODUCTION

The embryonic pharyngeal apparatus (PhAp) is a developmental system that provides progenitors and instructions to multiple organs and tissues, including but not limited to the craniofacial and mediastinic muscles and bones, most of the heart, and glands such as thymus, parathyroids and thyroid. Developmental anomalies of the PhAp underlie numerous birth defects, highlighting its developmental and genetic complexity. A textbook example of PhAp maldevelopment is DiGeorge syndrome, the most common genetic cause of which is a heterozygous deletion of a chromosomal region within 22q11.2 (in which case the clinical presentation is more complex and is designated as 22q11.2 deletion syndrome), and it can also be caused by point mutations of the *TBX1* gene (Haddad et al., 2019; Paylor et al., 2006; Xu et al., 2014; Yagi et al., 2003; Zweier et al., 2007).

The development of the PhAp depends upon the contribution of tissues derived from all three germ layers: surface ectoderm, pharyngeal endoderm, neural crest-derived cells (NCCs) and the cardiopharyngeal mesoderm (CPM). The latter contributes to a broad range of tissues and structures within the mediastinum and face and neck (Adachi et al., 2020). In the mouse, the CPM is well represented by the expression domains of the *Tbx1^Cre^* and the *Mef2c-AHF-Cre* drivers (Adachi et al., 2020; Huynh et al., 2007; Verzi et al., 2005). PhAp lineages have distinct origins and transcriptional profiles (Swedlund and Lescroart, 2020; Wang et al., 2019), develop in close proximity or direct contact with each other, and their regionalization within the PhAp is mostly conserved across vertebrate evolution (Graham, 2001). However, the molecular code that governs regionalization has not been dissected in detail, although interactions between lineages are the subject of intense research (Calmont et al., 2009; Huang et al., 1998; Kodo et al., 2017; Mao et al., 2021; Sato et al., 2011; Shone and Graham, 2014; Warkala et al., 2020).

Loss of function of the *Tbx1* gene in the mouse has profound and broad effects on the development of the PhAp (Jerome and Papaioannou, 2001; Lindsay et al., 2001; Merscher et al., 2001), and affects the expression of thousands of genes (Fulcoli et al., 2016; Ivins et al., 2005; Liao et al., 2008; Pane et al., 2012), making it difficult to identify the effectors/targets that are critical for specific developmental functions. Phenotypic rescue strategies represent an alternative approach to focus on genes associated with phenotypic improvement.

In a search for drugs that rebalance *Tbx1* haploinsufficiency, we showed that high doses of vitamin B12 (vB12) rescued part of the mutant phenotype *in vivo* (Lania et al., 2016). Here, we show that the rescuing capacity of the drug extends to CPM-derived structures, such as the cardiac outflow tract and craniofacial muscles. Then, as a proof of principle of the usefulness of phenotypic rescue to provide insights into pathogenetic mechanisms, we leveraged vB12 treatment to identify genes and pathways that are critical for the expressivity of the rescued phenotype.

This exposed a novel *Tbx1* mutant phenotype through the identification of a SLUG-positive subpopulation of CPM cells. Specifically, we found that in *Tbx1* homozygous mutants, SLUG+ cells were segregated from the NCCs rather than intermingled with them, suggesting a cell sorting defect. This abnormality was also evident in *Tbx1* heterozygous mutants, albeit at a reduced expressivity. Thus, in the PhAp, TBX1 dosage is important, cell autonomously and non-autonomously, for the regionalization of cell lineages. We propose that this TBX1-dependent function is part of the pathogenetic mechanism leading to severe abnormalities of the PhAp in the mouse mutants as well as in DiGeorge syndrome.

## RESULTS

### Vitamin B12 reduces the severity of the intracardiac and craniofacial phenotypes in a hypomorphic Tbx1 mutant model

High dosage of vB12 reduced the penetrance of the aortic arch phenotype and rebalanced *Tbx1* expression in haploinsufficient mice (Lania et al., 2016). However, *Tbx1*^+/-^ embryos rarely show second heart field (SHF)-related abnormalities such as outflow tract defects, which are commonly found in embryos that express low levels of *Tbx1* (Liao et al., 2004; Zhang and Baldini, 2008) or in *Tbx1*^-/-^ embryos (Jerome and Papaioannou, 2001; Lindsay et al., 2001; Merscher et al., 2001). We asked wether vB12 treatment, could modify the SHF-dependent phenotype on a *Tbx1* reduced-dosage model. To this end, we exploited a hypomorphic *Tbx1* allele (*Tbx1^neo2^*) (Zhang et al., 2006) that has a *lox*P-flanked neomycin resistance gene inserted into an intron. *Tbx1^neo2/-^* embryos exhibit heart defects similar to but less severe than null embryos (Zhang and Baldini, 2008). We crossed *Tbx1^neo2/+^* and *Tbx1*^+/-^ mice, and injected pregnant females daily from embryonic day (E) 7.5 to E11.5 with vB12 (intraperitoneal, i.p. injection, 20mg/Kg/day) or vehicle (PBS, controls). Embryos were harvested and dissected at E15.5 and E18.5. Table 1 summarizes the phenotyping results. We examined 17 *Tbx1^neo2/-^* embryos at E18.5 (8 controls and 9 treated with vB12).

**Table1.** Summary of heart morphology phenotyping data

| Genotype | n | Treatment | Normal | VSD | OvAo | DORV | incPTA | PTA |
| --- | --- | --- | --- | --- | --- | --- | --- | --- |
| E18.5 |  |  |  |  |  |  |  |  |
| *Tbx1*<sup>neo2/-</sup> | 8 | PBS | 0 | 8(100%) | 0 | 0 | 0 | 8(100%) |
| *Tbx1*<sup>neo2/-</sup> | 9 | vB12 | 1(11%) | 8(100%) | 2(25%) | 2(22%) | 4(44%) | 2(22%) |
| E15.5 |  |  |  |  |  |  |  |  |
| *Tbx1*<sup>neo2/-</sup> | 5 | PBS | 0 | 5(100%) | 0 | 0 | 0 | 5(100%) |
| *Tbx1*<sup>neo2/-</sup> | 3 | vB12 | 0 | 3(100%) | 3(100%) | 0 | 0 | 0 |

*DORV: Double outlet right ventricle; OvAo: overriding aorta; incPTA: incomplete PTA; PTA: persistent truncus arteriosus; VSD: ventricular septal defects.*

All control *Tbx1^neo2/-^* embryos had persistent truncus arteriosus (PTA) and ventricular septal defects (VSD) (Fig. 1A-A’). In contrast, only 2 of the 9 B12-treated embryos exhibited typical PTA (Fig. 1B), while 4 had an incomplete PTA, in which there was an unseptated valve but a distal separation of the aorta and pulmonary trunk (Fig. 1B’), and 2 had double outlet right ventricle (DORV) (Fig. 1-B”). All embryos had a VSD, with the exception of one embryo, which had an apparently normal heart (Fig. 1C’, Supplementary Fig. 1). In addition, we have examined histologically a set of 5 control and 3 vB12-treated *Tbx1^neo2/-^* embryos at E15.5. All control embryos had VSD and PTA, while the B12-treated embryos had VSD and overriding of the aorta, but no PTA (Table 1 and Supplementary Fig. 2). Reduced dosage of *Tbx1* causes specific craniofacial muscle anomalies (Adachi et al., 2020; Dastjerdi et al., 2007; Kelly et al., 2004). We tested a set of 5 controls and 3 B12-treated *Tbx1^neo2/-^* embryos at E15.5 and scored the craniofacial muscle phenotype (Fig. 2 and Supplementary Fig. 3). Results showed that vB12 treatment reduced the severity of anomalies of the muscles originating from the 1st pharyngeal arch (PA); bilateral defects of the anterior digastric muscles in *Tbx1^neo2/-^* embryos reduced from 60% to 33% after vB12 treatment. Defects of 2nd PA-derived branchiomeric muscles were also rescued by vB12 treatment (Fig. 2 and Supplementary Table 1). However, it did not have any effect on muscles derived from more posterior PAs (Supplementary Fig. 3 and Supplementary Table 1).

**Figure 1.**
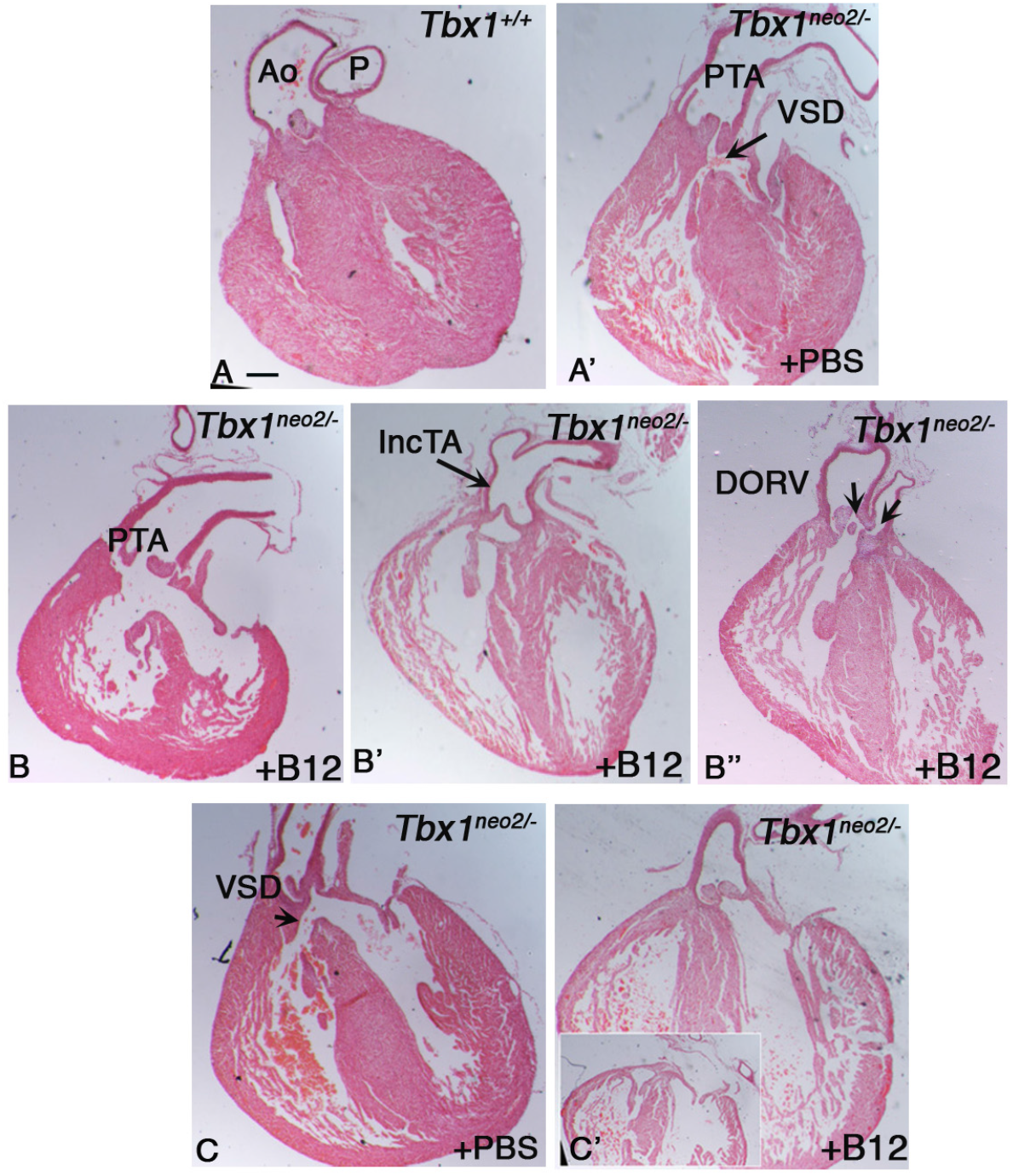
Vitamin B12 ameliorates cardiac outflow tract anomalies observed in Tbx1^neo2/-^ mouse embryos. *Representative images of histological sections (coronal) of heart from Tbx1^+/+^ and Tbx1^neo2/-^ embryos at E18.5, treated with vB12 or PBS. A) Histological sections of hearts from Tbx1^+/+^ +PBS and B) Tbx1^neo2/-^ +PBS control embryos. Aorta (Ao) and pulmonary trunk (P) are separated. C.D.E) Histological sections of hearts from Tbx1^neo2/-^ + vB12 embryos; PTA: Persistent truncus arteriosus; IncTA: incomplete truncus arteriosus; DORV: double outlet right ventricle. F) Histological sections of hearts from Tbx1^neo2/-^ +PBS and G) vB12 treated embryos; VSD: ventricular septal defect. Scale bar: 200 μm*

**Figure 2.**
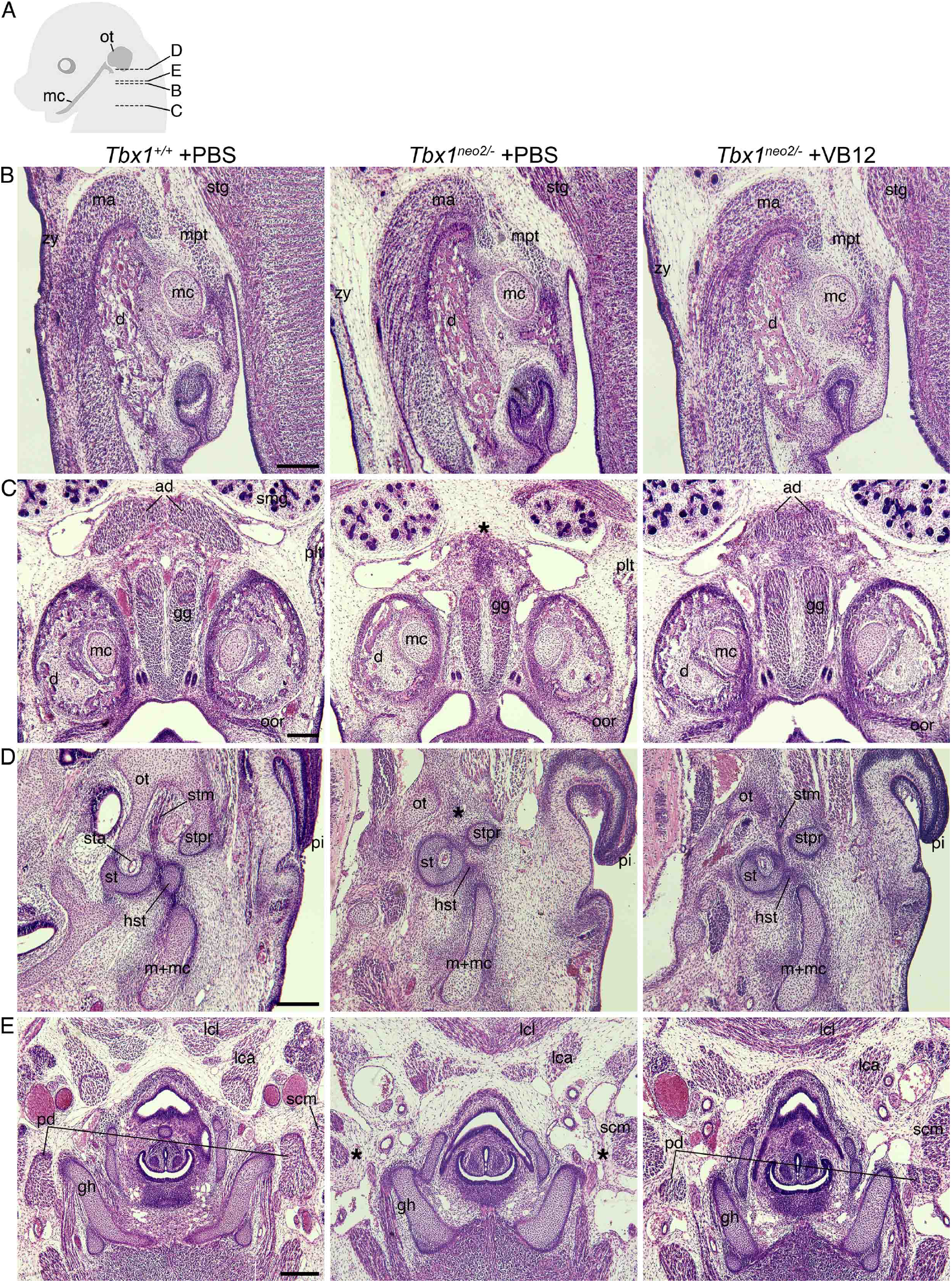
Vitamin B12 treatment partially rescues the branchiomeric muscle phenotype in *Tbx1^neo2/-^* embryos. *Histological sections of Tbx1^+/+^ and Tbx1^neo2/-^ embryos at E15.5, stained with hematoxylin and eosin. A) A diagram showing the section levels. B) Middle and C) ventral jaw muscles derived from 1st pharyngeal arch. D) Otic and E) hyoid muscles derived from 2nd pharyngeal arch. The asterisks indicate missing muscles that are rescued by vB12, but not by PBS treatment, in this particular set of embryos (anterior digastric, posterior digastric, and stapedius muscles). For a complete list of results, see Supplementary Table 1.* *Abbrevations: <u>ad, anterior digastric muscle</u>; d, dentary; gg, genioglossus muscle; gh, greater horn of hyoid bone; hst, head of stapes; lca, longs capitis muscle; lcl, longs coli muscle; ma, masseter muscle; mc, Meckel’s cartilage; m+mc, malleus and Meckel’s cartilage; mpt, medial pterygoid muscle; oor, orbicularis oris muscle; ot, otic capsule; <u>pd, posterior digastric muscle</u>; pi, pinna; plt, platysma muscle; scm, sternocleidomastoid muscle; smg, submandibular gland; st, stapes; sta, stapedial artery; stg, styloglossus muscle; <u>stm, stapedius muscle</u>; stpr, styloid process; zy, zygomaticus muscle.* *Scale bars: 200 μm.*

### Identification of rescued genes after B12 treatment in vivo

In order to evaluate the effect of vB12 treatment on embryo tissue transcription, we performed RNA-seq on whole E9.5 mouse embryos (21 somite stage) after treatment with vB12 or vehicle (PBS) during pregnancy (i.p. 20mg/Kg/day at E7.5, E8.5, and E9.5). We analyzed the results from PBS-treated WT (n=3), PBS-treated *Tbx1^+/-^* (n=3), and vB12-treated *Tbx1^+/-^* (n=2) embryos, where each embryo was sequenced independently and each dataset was treated as a biological replicate. Comparing *Tbx1*^+/-^ *vs Tbx1^+/+^* embryos, we found a total of 1409 differentially expressed genes (DEGs) (fold change cut off of > 1.2, and posterior probability, PP > 0.95) of which 851 (60.4%) up-regulated and 558 (39.6%) were down-regulated (Fig. 3A and Supplementary Tab. 2). Gene ontology analyses of the 851 up-regulated genes revealed an enrichment in genes involved in oxidative phosphorylation and other metabolic processes, while among the 558 down-regulated genes, there was enrichment of genes involved in morphogenesis and developmental processes (Table 2). Comparing vB12-treated *Tbx1^+/-^* embryos with PBS-treated *Tbx1^+/-^* embryos, we found a total of 3954 DEGs, of which 1862 (47%) were up-regulated and 2092 (53%) where down-regulated (Fig. 3B and Supplementary Tab. 2). Gene ontology analyses revealed that the up-regulated genes were enriched for genes involved in RNA processing, while down-regulated genes were enriched for genes involved in developmental processes (Table 3). Details of the gene ontology analyses are shown in Supplementary Table 3.

**Figure 3.**
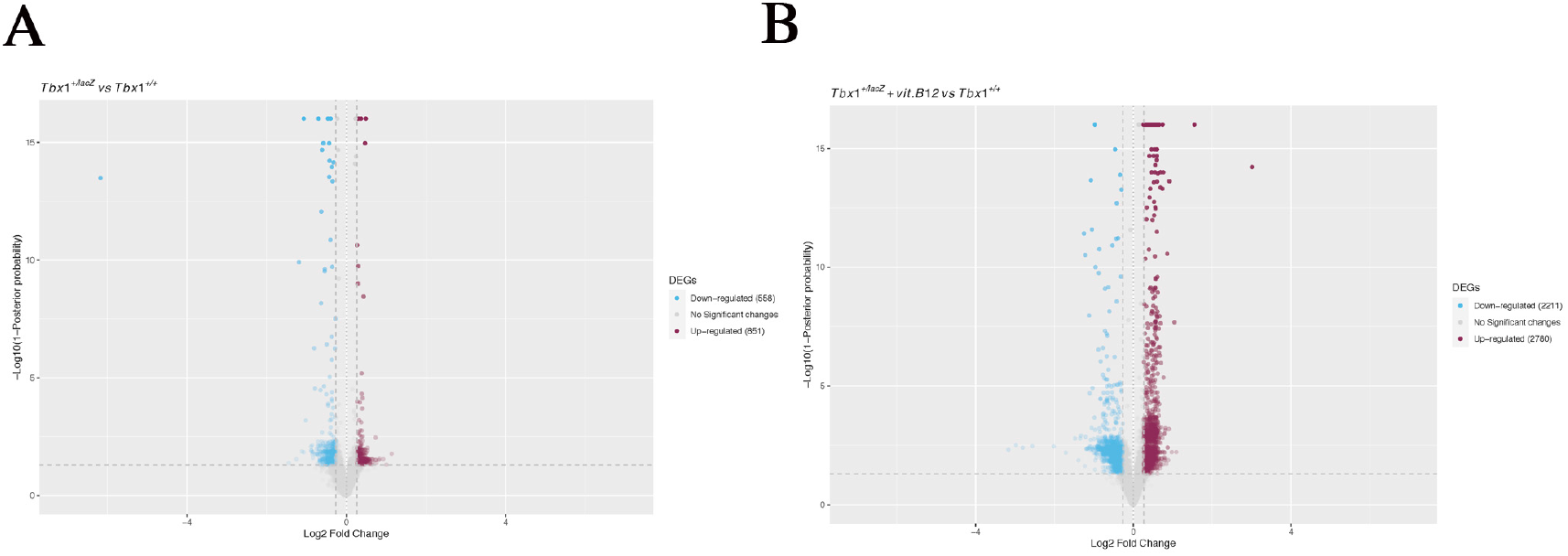
Tbx1 gene haploinsufficiency alters the expression of 1409 genes, while Vitamin B12 treatment in heterozygous background induces dysregulation of 3954 genes. *A) Volcano plot of significantly up regulated or down regulated genes in Tbx1^+/-^ whole embryos compared to Tbx1^+/+^ embryos. Blue dots represent down regulated genes, red dots represent up regulated genes. B) Volcano plot of significantly up regulated or down regulated genes in Tbx1^+/-^ + vB12 embryos compared to controls. Blue dots represent down regulated genes, red dots represents up regulated genes.*

**Table 2.** Gene ontology analysis of down regulated and up regulated genes of DEGs from comparison Tbx1^+/-^ vs Tbx1^+/+^

| 851 Up regulated genes |  |  | 558 Down regulated genes |  |  |
| --- | --- | --- | --- | --- | --- |
| term_name | term_id | Adjusted<br>_p_value | term_name | term_id | adjusted<br>_p_value |
| peptide biosynthetic process | GO:0043043 | 3,46E-20 | anatomical structure morphogenesis | GO:0009653 | 8,51E-14 |
| peptide metabolic process | GO:0006518 | 8,08E-20 | cellular component organization | GO:0016043 | 1,58E-11 |
| oxidative phosphorylation | GO:0006119 | 4,84E-16 | system development | GO:0048731 | 5,65E-11 |
| amide biosynthetic process | GO:0043604 | 6,06E-16 | developmental process | GO:0032502 | 8,41E-11 |
| mitochondrial respiratory chain complex assembly | GO:0033108 | 7,71E-15 | multicellular organism development | GO:0007275 | 8,68E-11 |
| electron transport chain | GO:0022900 | 1,04E-13 | anatomical structure development | GO:0048856 | 9,92E-11 |
| mitochondrial respiratory chain complex I assembly | GO:0032981 | 5,89E-13 | cell development | GO:0048468 | 1,18E-10 |
| NADH dehydrogenase complex assembly | GO:0010257 | 5,89E-13 | heart development | GO:0007507 | 2,97E-10 |
| cellular amide metabolic process | GO:0043603 | 1,15E-12 | cellular component organization or biogenesis | GO:0071840 | 3,08E-10 |
| ATP synthesis coupled electron transport | GO:0042773 | 1,65E-11 | nervous system development | GO:0007399 | 1,00E-09 |
| mitochondrial ATP synthesis coupled electron transport | GO:0042775 | 3,52E-11 | embryo development | GO:0009790 | 2,40E-09 |

**Table 3.** Gene ontology analysis of down regulated and up regulated genes of DEGs from comparison Tbx1^+/-^ + B12 vs Tbx1^+/-^ +PBS

| 2780 Up regulated genes |  |  | 2211 Down regulated genes |  |  |
| --- | --- | --- | --- | --- | --- |
| term_name | term_id | adjusted<br>p value | term_name | term_id | Adjusted<br>_p_value |
| ribonucleoprotein complex biogenesis | GO:0022613 | 1,12E-23 | nervous system development | GO:0007399 | 7,91E-56 |
| ribosome biogenesis | GO:0042254 | 1,53E-23 | multicellular organism development | GO:0007275 | 8,63E-47 |
| cellular nitrogen compound metabolic process | GO:0034641 | 5,27E-20 | system development | GO:0048731 | 1,46E-46 |
| RNA processing | GO:0006396 | 5,56E-19 | regulation of transcription by RNA polymerase II | GO:0006357 | 2,77E-42 |
| ncRNA processing | GO:0034470 | 5,74E-19 | transcription by RNA polymerase II | GO:0006366 | 1,22E-40 |
| ncRNA metabolic process | GO:0034660 | 1,88E-17 | neurogenesis | GO:0022008 | 1,92E-39 |
| rRNA processing | GO:0006364 | 3,72E-15 | regulation of cellular process | GO:0050794 | 2,90E-39 |
| rRNA metabolic process | GO:0016072 | 3,95E-14 | cellular developmental process | GO:0048869 | 1,57E-38 |
| ribosomal large subunit biogenesis | GO:0042273 | 9,62E-13 | anatomical structure development | GO:0048856 | 1,93E-38 |
| heterocycle metabolic process | GO:0046483 | 1,05E-11 | cell differentiation | GO:0030154 | 2,21E-38 |
| cellular metabolic process | GO:0044237 | 1,73E-11 | anatomical structure morphogenesis | GO:0009653 | 1,66E-37 |
| cellular aromatic compound metabolic process | GO:0006725 | 6,17E-11 | developmental process | GO:0032502 | 2,02E-37 |

The intersection of the two groups of DEGs identified 468 shared genes (Fig. 4A), which is significantly higher than that expected by chance (P=1.4×10^-7^, hypergeometric test). Of these, 344 changed their expression in opposite directions in the two groups, i.e. the mutation changed expression in one direction while vB12 treatment rebalanced it (Fig. 4B, genes listed in Supplementary Tab. 2); we define these genes as rescued by vB12. In the left column of the heat map shown in Fig. 4B are represented genes dysregulated by the mutation; the right column shows their expression after vB12 treatment (compared to WT). Thus, the dark color in the right column indicates genes that were expressed at or near WT level after treatment. Of these 344 genes, 85 (24.7%, shown in green) were down-regulated, and 259 (75.3%, shown in red) were up-regulated relative to WT (Fig. 4B, left column).

**Figure 4.**
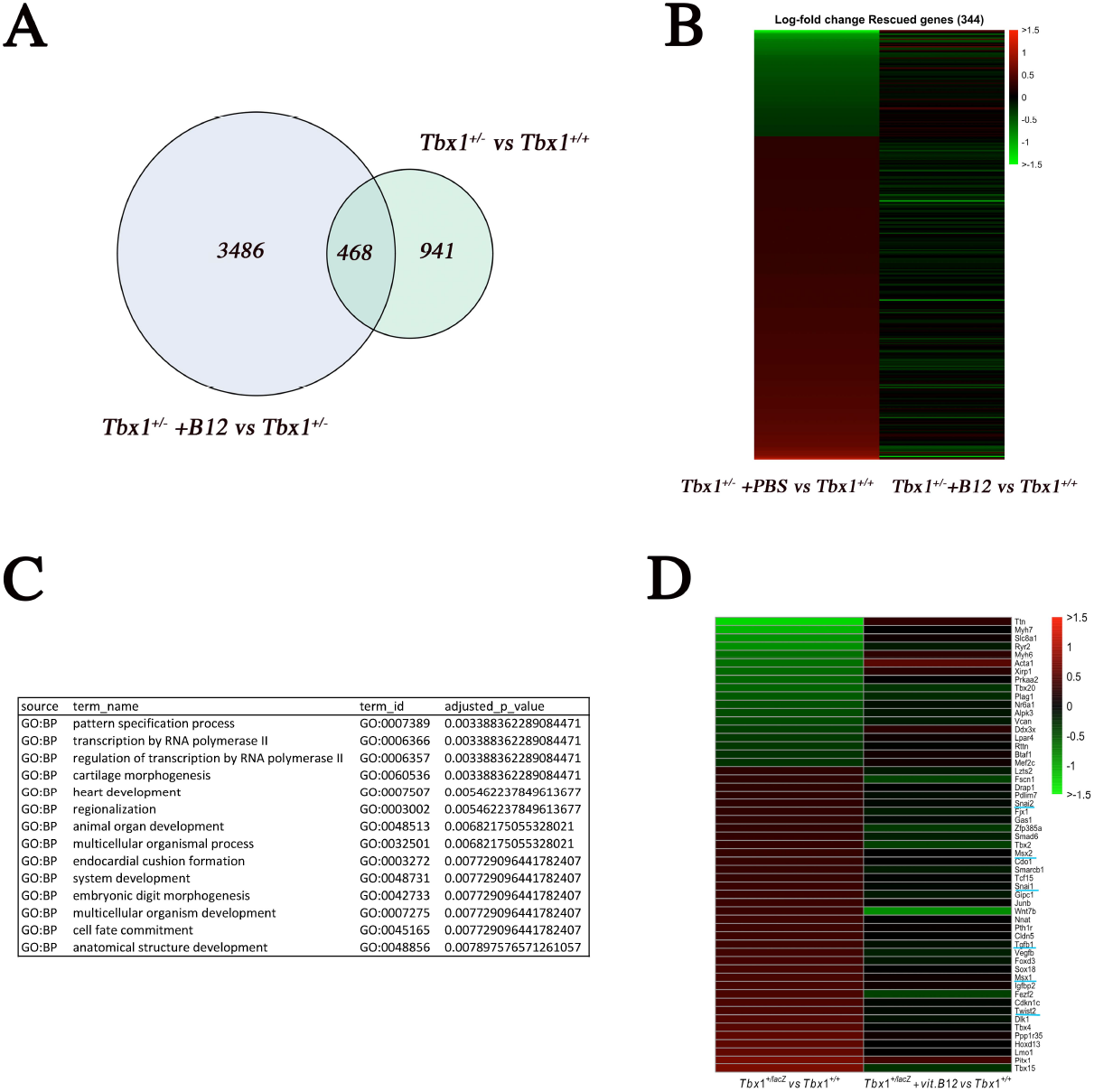
Vitamin B12 rescues the expression of 344 genes involved in gene regulation and heart development. *A) Venn diagram plot representing the intersection of two groups of differentially expressed genes (Tbx1^+/^ vs. Tbx1^+/+^, and Tbx1^+/^ +vB12 vs. Tbx1^+/^ +PBS). B) Heat maps of rescued genes in Tbx1^+/^ embryos (treated with vB12 or PBS) based on their fold change relative to WT. C) gene ontology analysis by the gProfiler tool. D) Heat map of 55genes selected because known to be expressed in the pharyngeal apparatus.*

We then applied a hypergeometric test to ask whether rescue of gene expression imbalances by vB12 could have occurred by chance, given the high number of genes affected by the treatment. Interestingly, we found that the rescue of the 259 up-regulated genes was extremely significant (P<<10^-10^), while the rescue of the 85 downregulated genes was borderline with a chance event (P=0.052). In addition, the number of genes that were further dysregulated by vB12 (468-344=124), which were almost equally distributed among up- and down-regulated (59 and 65, respectively) were not significantly different to that expected from a chance event. Thus, the most significant rescue effect of vB12 was on genes that were up-regulated by *Tbx1* heterozygosity and down regulated by vB12 treatment.

Gene ontology of the 344 rescued genes showed enrichment of Heart Development genes (P=0.005) and Transcription Regulator genes (P=0.003) (Supplementary Table 3).

### SLUG identifies a mesodermal population partially overlapping with but distinct from the TBX1+, ISL1+, and Mef2c-AHF-Cre+ populations

Among the rescued genes, we identified 55 genes known to be expressed in the cardiopharyngeal mesoderm (CPM), or its derivatives, and to a lesser extent, in other tissues of the PhAp (Fig. 4D and Supplemetary Tab. 2). Among these, we noted a set of genes known to be involved in the Tgfβ1 pathway. Specifically, *Snai1, Snai2, Twist2, Msx1,* and *Tgfb1* were all up-regulated in *Tbx1^+/-^* mutant embryos and down regulated by vB12 treatment (blue underlined in Fig. 4D). We selected *Snai2*, encoding a transcriptional repressor also known as SLUG, which has not been previously associated with TBX1 biology. We performed immunofluoresence (IF) with an anti-SLUG antibody to determine its expression relative to markers of the CPM in E9.5 WT embryos. The anti-SLUG and anti-TBX1 antibodies are raised in the same species, therefore we used them in sequence on the same sections; similar results were obtained by inverting the order of the antibodies. As shown in Fig. 5A-B, at all section levels considered, there was a very similar distribution of the two proteins, with notable exceptions. Specifically, in the 1st PA, both proteins were present in the core mesoderm, but SLUG+ cells were fewer in the core and there were more of them scattered in the body of the PA (arrowhead in B); TBX1+ cells were more evident in the proximal region of the arch (Fig. 5A-B). In the 2nd PA, the SLUG domain extended more distally towards the OFT, compared to the TBX1 domain (Fig. 5A’-B’). At the other two levels analysed, namely, posterior to the OFT (Fig. 5A”-B”) and immediately anterior to the inflow tract (IFT) (Fig. 5A’”-B’”), the distribution of the two proteins was very similar in the lateral aspects of the dorsal pericardial and splanchnic mesoderm.

**Figure 5.**
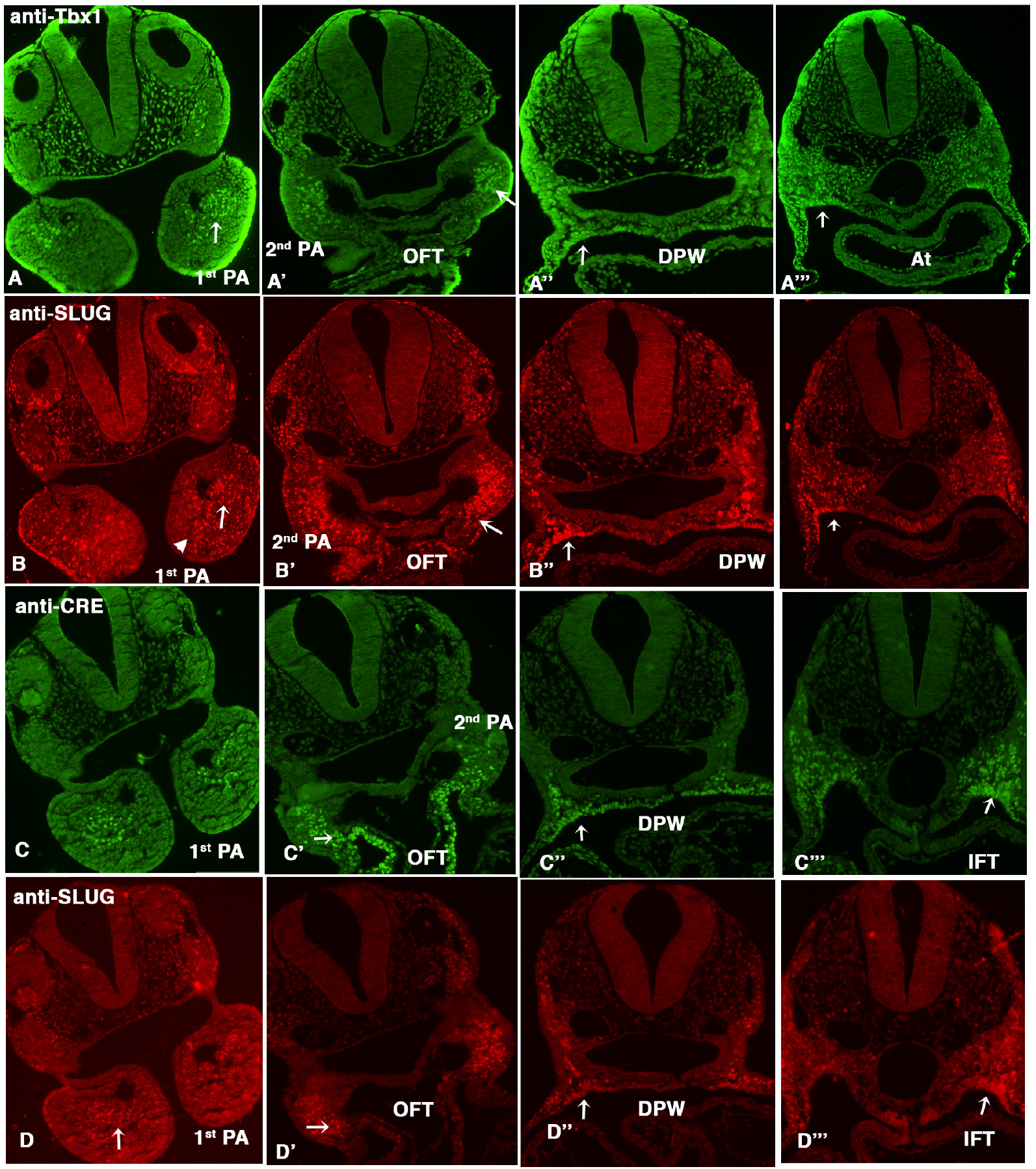
SLUG is expressed in the mesoderm and partially overlaps with TBX1 and Mef2c-AHF expression. *Immunofluorescence on transverse sections of E9.5 embryos. A and B, comparison of TBX1 and SLUG expression A) anti TBX1 antibody, B) anti SLUG antibody on the same sections; C and D, comparison of the expression patterns of Cre driven by Mef2c-AHF-Cre, and SLUG. C) anti-Cre and D) anti SLUG. For both comparisons, we have used sequential staining because the antibodies were raised in the same species. The arrows indicate the expression of Tbx1 or CRE or SLUG in core mesoderm region in 1^st^ PA, in the distal mesoderm in 2^nd^ PA, in dorsal pericardial wall and in second heart field.* *PA: Pharyngeal arch; DPW: dorsal pericardial wall; SHF: Second heart field, IFT: inflow tract; OFT: outflow tract.* *Scale bar: 100 μm*

Next, we compared the expression of SLUG to that of the Mef2c-AHF enhancer using an anti-CRE antibody on sections of Mef2c-AHF-Cre embryos (Verzi et al., 2005). Also in this case, the expression patterns of the two proteins were very similar (Fig. 5C-D), except for two substantial differences; in the 2nd PA, Mef2c-AHF-Cre was expressed more distally towards the OFT, including the myocardial layer of the OFT, (Fig. 5C’-D’), more extensively in the dorsal pericardial wall (Fig. 5C”), and more extensively in the splanchnic mesoderm of the posterior region of the embryonic pharynx (Fig. 5C’”-D’”).

We then compared SLUG expression to that of ISL1, which is expressed throughout the CPM (Fig. 6). The two markers had a very similar, mostly overlapping expression in the 1st pharyngeal arch (Fig. 6A-B) of WT embryos. In the 2nd arch, SLUG was more highly expressed in the distal portion of the arch, where it overlapped with ISL1, whereas in the proximal portion, there was a high expression of ISL1 but not of SLUG. Double labeled cells were detected in a medio-lateral region of the dorsal pericardial wall, were ISL1 expression was more extensive (Fig. 6A”-B”); in addition, the more posterior expression of ISL1 in the splanchnic mesoderm (arrow in Fig. 6B”) is similar to the TBX1 expression domain (Fig. 6C-D).

**Figure 6.**
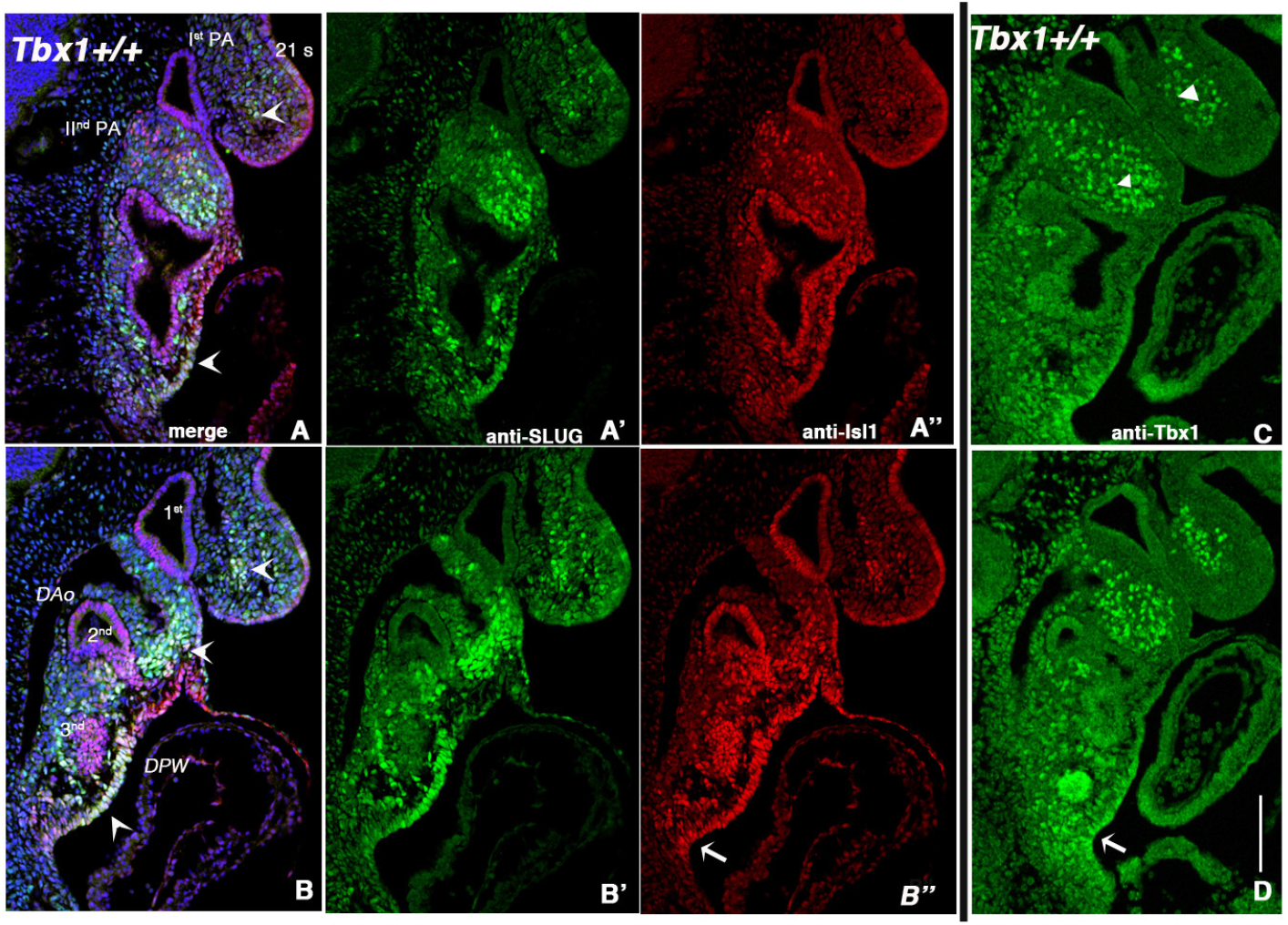
SLUG and ISL1 have a similar pattern of expression with exceptions in the proximal 2nd PA and pSHF. (*A-B) Double immunofluorescence with anti-SLUG and anti-ISL1 on sagittal sections of WT E9.5 embryos. Two representative images from lateral (A) to medial (B). A’-B’) anti-SLUG; A”-B”) anti-ISL1. Arrows indicate regions where ISL1 expression is more extensive than SLUG (compare A’ with A”, and B’ and B”. In C) and D) are shown similar sections but immunostained with an anti-TBX1 antibody. Note a similar expression of TBX1 and ISL1 at this level (arrows). Note also the difference in the expression in the 1st PA (arrowhead) compared to both ISL1 and SLUG. PA: pharyngeal arch; DPW: dorsal pericardial wall; DAo: dorsal aorta. Scale bar: 200 μm*

Because of the scattered expression of SLUG in the body of the 1st PA, which is heavily populated by NCCs, we co-stained E9.5 embryos (WT) with TFAP2A, which labels migrating NCCs at this stage (as well as ectodermal cells). With very few exceptions, we did not observe double labeled cells in the entire embryo (examples in Fig. 7A, C, E), but we observed extensive intermingling of SLUG+ and TFAP2A+ cells. Thus, SLUG is expressed in mesodermal cells, mainly in the distal 2nd PA, in the medio-lateral dorsal pericardial wall, in the posterior lateral splancnic mesoderm and, to a minor extent, in the 1st PA.

**Figure 7.**
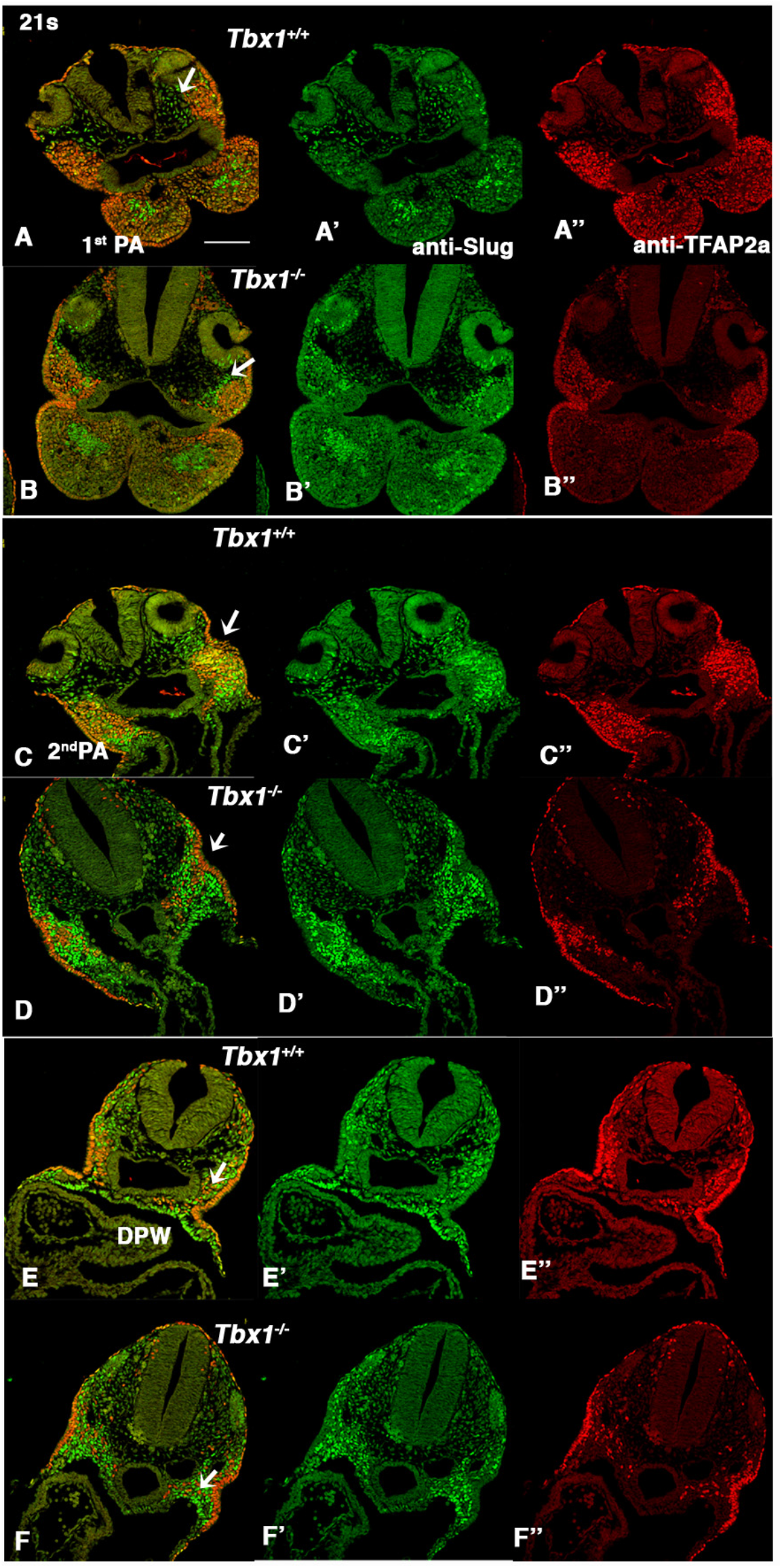
TFAP2A and SLUG highlights regionalization defects in *Tbx1^-/-^* embryos. *Representative images of double immunofluorescence anti TFAP2 (red) and SLUG (green) transverse sections of Tbx1^+/+^ (A,C,E) and Tbx1^-/-^ (B,D,F) E9.5 embryos at the level of the 1st PA (A-B), Ilnd PA/OFT (C-D), and dorsal pericardial wall (DWP) between the OFT and IFT (E-F). In general, there is minimal or no overlap between the two markers. In A-B note the different relative distribution of SLUG+ and TFAP2A+ cells in the 1st PA (arrows). In C-D, cells of the two lineages are intermingled in the WT 2nd PA but segregated in the mutant (arrows); in E-F, note the expansion of SLUG expression in the mutant (arrows). Scale bar: 200 μm*

### Tbx1 mutants have lineage regionalization abnormalities

We next investigated whether *Tbx1* loss of function affected the SLUG+ population. We first examined this population in comparison to NCCs (TFAP2+; TBX1-negative). At the level of the 1st PA, we found a striking pattern in which SLUG+ cells in *Tbx1^-/-^* embryos were tightly grouped in the core mesoderm forming a large area surrounded by, but not mixed with TFAP2A+ cells (Fig. 7B), while in WT embryos the two cell types were intermingled (Fig. 7A). A similar pattern was evident in the head mesoderm/proximal 1^st^ PA (Fig. 7A-B, arrows). At the level of the 2^nd^ PA, which is severely hypoplastic in *Tbx1*^-/-^ embryos, the mixing of the two populations was substantially reduced, although here the TFAP2+ population appeared smaller than in WT (Fig. 7C-D, arrows). More posteriorly (caudal to the OFT) this segregation phenotype was not apparent, of note is a relative expansion of the SLUG+ population at this level in the splanchnic mesoderm of the mutant embryo (Fig. 7E-F, arrows).

### Tbx1 mutants have lineage regionalization abnormalities

We next investigated whether *Tbx1* loss of function affected the SLUG+ population. We first examined this population in comparison to NCCs (TFAP2+; TBX1-negative). At the level of the 1st PA, we found a striking pattern in which SLUG+ cells in *Tbx1^-/-^* embryos were tightly grouped in the core mesoderm forming a large area surrounded by, but not mixed with TFAP2A+ cells (Fig. 7B), while in WT embryos the two cell types were intermingled (Fig. 7A). A similar pattern was evident in the head mesoderm/proximal 1^st^ PA (Fig. 7A-B, arrows). At the level of the 2^nd^ PA, which is severely hypoplastic in *Tbx1*^-/-^ embryos, the mixing of the two populations was substantially reduced, although here the TFAP2+ population appeared smaller than in WT (Fig. 7C-D, arrows). More posteriorly (caudal to the OFT) this segregation phenotype was not apparent, of note is a relative expansion of the SLUG+ population at this level in the splanchnic mesoderm of the mutant embryo (Fig. 7E-F, arrows).

We next examined the distribution of SLUG+ cells compared to ISL1+ cells in *Tbx1^-/-^* embryos. In the 1st PA of *Tbx1*^-/-^ embryos, and in contrast to WT embryos, we observed a large, well defined cluster of SLUG+ cells that were mostly ISL1+ in the core mesoderm and appeared to extend posteriorly, as if it resulted from merging with the core of the 2nd PA, which is severely hypoplastic in these mutants (arrow in Fig. 8A-B). In a more medial sagittal plane, the aggregate is also clearly visible (Fig. 8C-D).

**Figure 8.**
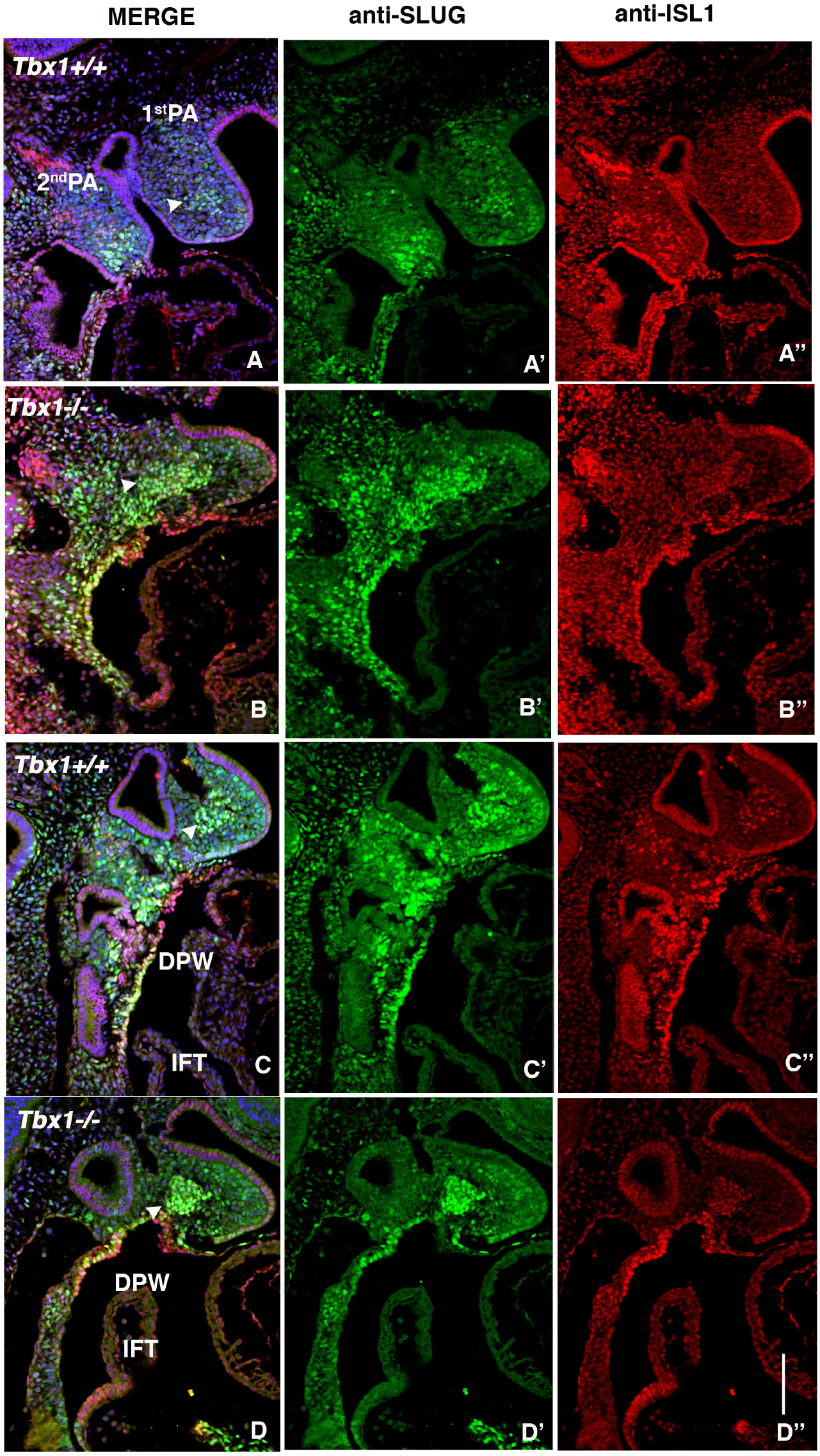
The core mesoderm of the 1st PA of *Tbx1^-/-^* embryos is populated by SLUG+;ISL1+ cells. *Double immunofluorescence using anti-SLUG (green) and anti-ISL1 (red) on medial sagittal sections (A,B) and on transverse sections (C,D) of 1st PA from Tbx1^+/+^ (A,C) and Tbx1^-/-^ (B,D) embryos. Note the co-localization of ISL1 and SLUG in the core mesoderm of 1st PA in Tbx1^-/-^. In the WT, this is much less evident (arrow in 1st PA). In addition, SLUG expression is extended in the splanchnic mesoderm of the Tbx1^-/-^ embryo (lower arrow in B, compare A” and B”). INF: Venous pole. Scale bar: 100 μm*

We next tested whether cells of the *Tbx1* genetic lineage are mislocalized relative to SLUG+ cells, in the absence of *Tbx1* function. To this end, we performed anti-SLUG and anti-GFP IF on *Tbx1^cre/-^;R26R^mT-mG^ (Tbx1* null) and *Tbx1^cre/+^;R26R^mT-mG^* (heterozygous, control) E9.5 embryos (*Tbx1^Cre^* is a null allele). Results showed that in control embryos, GFP+ cells (shown in red in Fig. 9A-B) were more prominent in the proximal and lateral aspects of the 1st PA relative to SLUG+ cells, with only a limited overlap. Moreover, the relationship between the markers was largely conserved in *Tbx1* null embryos (Fig. 9A-B), indicating that the SLUG+ aggregate in the 1st PA is mostly made of cells that did not activate *Tbx1* gene transcription.

**Figure 9.**
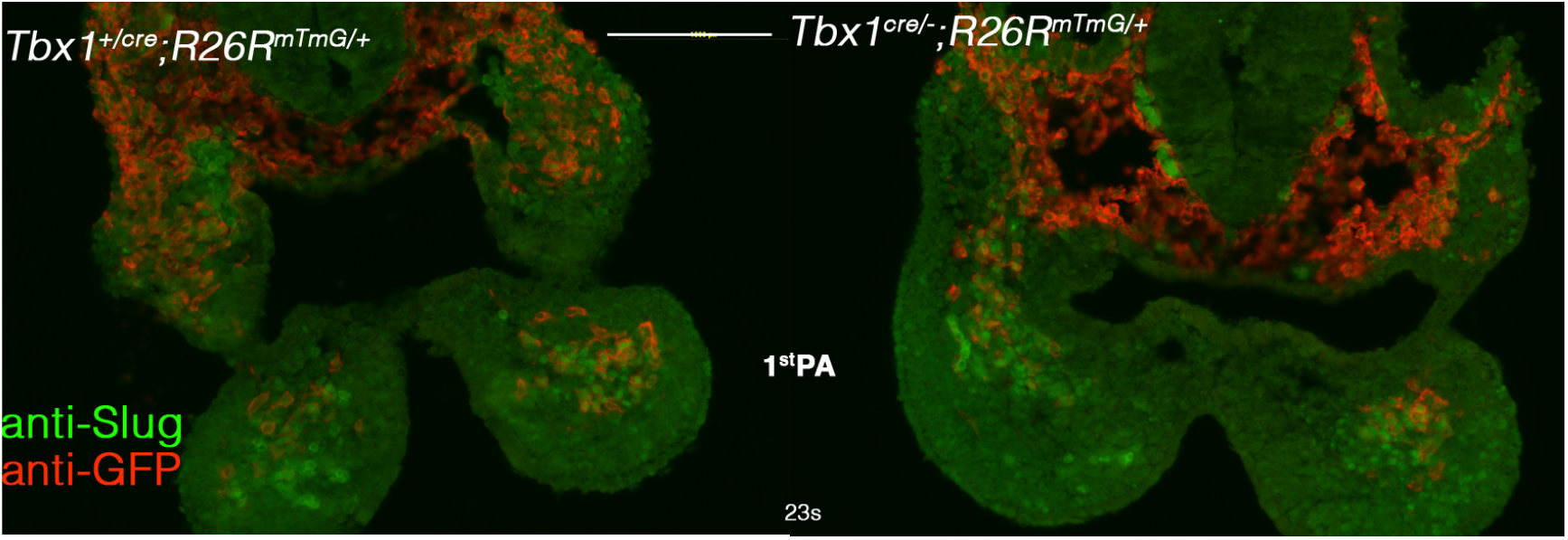
The positional relation between TBX1+ and SLUG+ cells is maintained in *Tbx1* null embryos. *Immunofluorescence using anti-GFP (red) and anti-SLUG (green) on transversal sections of Tbx1^Cre/+^;R26R^mTmG^ (functionally heterozygous, A) and Tbx1^Cre/-^;R26R^mTmG^ (null mutant, B) E9.5 embryos. Note that Tbx1-activating cells and their descendants (in red) localize prevalently in laterally in the core mesoderm and in the proximal region of the PA, in both cases. Scale bar: 100 μm*

To understand how the aggregation of SLUG+ cells in the 1st PA of *Tbx1*^-/-^ mutants arises, we examined earlier developmental stages: 11, 15, and 20 somites (st), immunostained with anti-SLUG and anti-ISL1 antibodies. In the WT embryo, at 11st the 1st PA is mostly populated by compacted mesoderm (ISL1+ and SLUG+) and a very limited non-mesodermal mesenchymal population (Supplementary Fig. 4A, A’). At this stage, the NCCs have not populated the arch in a substantial manner (Grenier et al., 2009). As NCCs populate the arch at 15st, they mostly surround the mesodermal core but a subpopulation invades the core, resulting in the dispersion of SLUG+ and ISL1+ cells within the arch mesenchyme (Supplementary Fig. 4B, B’). This process of dispersion continues at the 20st stage (Supplementary Fig. 4C, C’). In the *Tbx1*^-/-^ embryo, this process of dispersion does not occur at any stage (Supplementary Fig. 4D-F’), and as a result the SLUG+ and ISL1+ cells remain compacted. These observations suggest that in *Tbx1* mutants, NCCs fail to penetrate the pre-existing mesodermal core so that the two lineages remain segregated.

We next asked whether the segragation of SLUG+ cells may be explained by differential cell-cell adhesion mediated by cadherins. CDH2 (also known as N-Cadherin) is expressed in many mesodermal tissues and is involved in collective cell migration (review in (Alimperti and Andreadis, 2015)). We performed immunofluorescence with an anti-CDH2 anibody along with an anti-SLUG antibody and we found very low expression in the 1st PA of WT E9.5 embryos and higher expression in the Ilnd PA (Fig. 10B). However, in *Tbx1^-/-^* embryos, the compacted SLUG+ mesodermal core of the 1st PA was clearly CDH2+, well above the level of expression in the surrounding mesenchyme (Fig. 10C)

**Figure 10.**
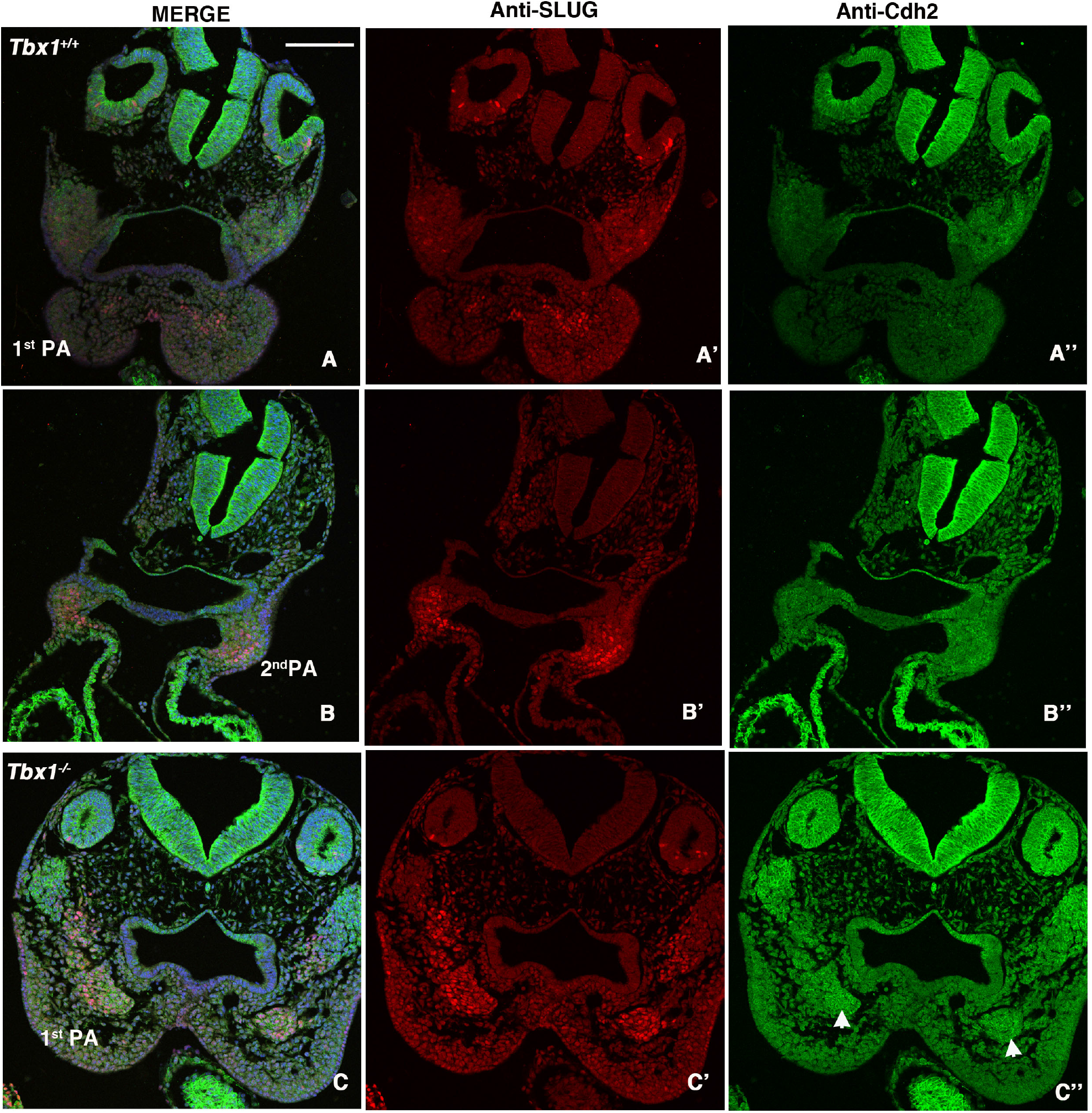
CDH2 (N-cadherin) is up-regulated in the 1st pharyngeal arch of *Tbx1^-/-^* E9.5 embryos. *Immunofluorescence of transverse sections of E9.5 embryos with the genotype indicated, double labelled with anti-SLUG (red) and anti CDH2 (green) antibodies. A-A”) transverse section at the level of the 1st PA, and B-B”) at the level of the 2nd PA and OFT of a WT embryo). C-C”) transverse section of a Tbx1-/- embryo at the level of the 1st PA, note the up regulation of CDH2 on the SLUG+ aggregate at the core of the arch (arrows). Scale bar: 25 μm*

In summary, our expression analysis indicates that TBX1 has cell autonomous and, perhaps more extensive non-cell autonomous functions in regulating the regionalization of cell lineages that are critical for the development of the PhAp.

### SLUG identifies a novel haploinsufficiency phenotype rescued by vB12

The results described above were obtained in *Tbx1*^-/-^ embryos, which exhibit significant anatomical anomalies, thus raising the question of whether some of the regionalization differences may be due to anatomical constraints. Therefore, we tested heterozygous mutants, which have no gross anatomical abnormalities (with the exception of hypoplasia of the 4th pharyngeal arch artery and parathyroids). As noted above, in the 1st PA of E9.5 WT embryos SLUG is expressed in a small number of cells of the core mesoderm and in scattered cells of the body of the arch (Fig. 11A). In *Tbx1^+/-^* embryos, SLUG+ cells were grouped in the core mesoderm (Fig 11A’, additional examples shown in Supplementary Fig. 5). A similar result was obtained using *Mef2c-AHF-Cre-driven* deletion of *Tbx1* in *Tbx1^flox/+^;Mef2c-AHF-Cre* embryos (Supplementary Fig. 6), indicating that this anomaly is dependent upon *Tbx1* haploinsufficiency in the mesoderm. This phenotype is reminiscent of but less severe than that noted in *Tbx1^-/-^* embryos (compare with Fig. 7A-B and Fig. 8C-D). Interestingly, vB12 treatment rescued this aggregation phenotype, re-establishing a cell distribution that was similar to the WT pattern (Fig. 11A”) in three independent experiments. Thus, even a 50% reduction of gene dosage is sufficient to generate defects in lineage regionalization, at least in the 1st PA, suggesting that these are unlikely to be explained by anatomical changes.

**Figure 11.**
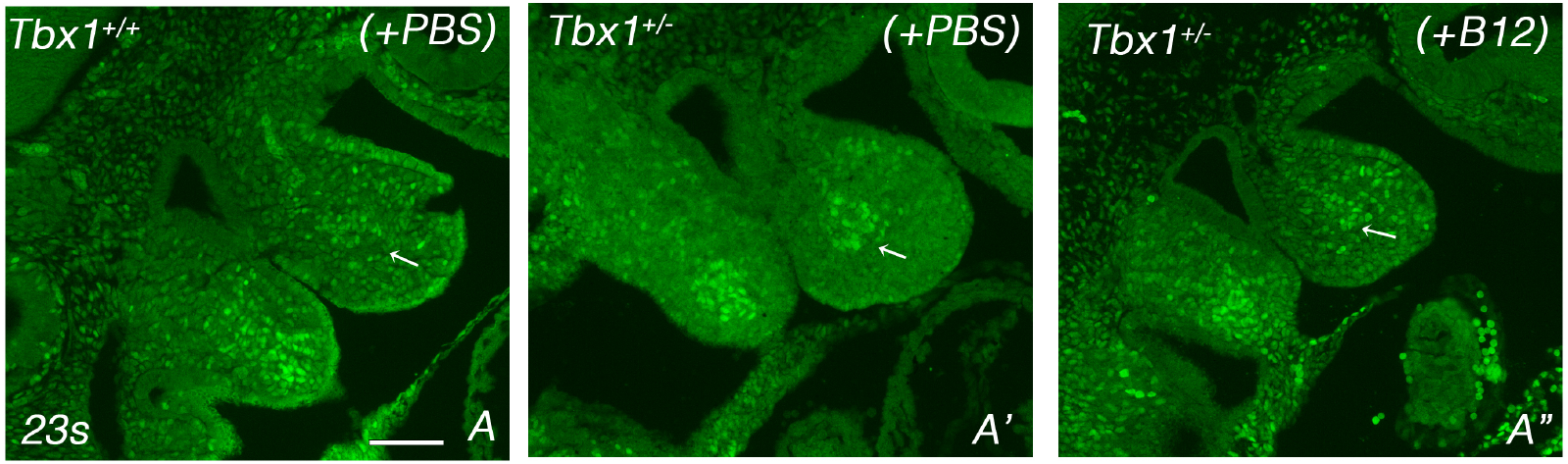
Tbx1 gene haploinsufficiency causes SLUG+ cell condensation in the 1st PA rescued by vitamin B12. *Images of immunofluorescence with an anti-SLUG antibody on WT+PBS (A), Tbx1^+/-^ +PBS (A’), and Tbx1^+/-^ +VIt. B12 (A”) E9.5 embryos. Medial-lateral sagittal sections. A cluster ofSLUG+ condensed cells in the core mesoderm is noticeable in A^r^, while in A” cells are more disperse, more similar to the WT pattern. Additional examples of the condensation phenotype are shown in Supplementary Fig. 5. Scale bar: 100 μm*

## DISCUSSION

Gene haploinsufficiency is a frequent cause of birth defects, and post-natal morbidity. Counter-balancing haploinsufficiency is possible but it is challenging in the clinical setting and may lack sufficient precision to rescue the full spectrum of phenotypic changes. Pharmacological rescue would be particularly useful in the clinics if it could be precisely targeted.

High doses of vB12 can partially rescue the 4th pharyngeal arch artery phenotype associated with *Tbx1* gene haploinsufficiency (Lania et al., 2016); in this study we tested additional phenotypic recoveries by crossing a hypomorphic allele, which combined with a null allele, causes a more complex phenotype than the one exhibited by heterozygous mutants. Indeed, we observed that vB12 treatment improved the septation process of the outflow tract in *Tbx1^neo2/^-* embryos. In addition, we found that branchiomeric muscle defects were also diminished after treatment.

Having shown that the treatment improves a range of phenotypic abnormalities, we sought to leverage this property to identify genes and pathways dysregulated by *Tbx1* haploinsufficiency and rebalanced by vB12, these genes may be critical for the pathogenesis of the rescued phenotypes. We found that in E9.5 *Tbx1*^+/-^ embryos, 24% of the genes dysregulated by *Tbx1* haploinsufficiency were rescued by vB12, and gene ontology analysis of the rescued genes revealed enrichment of genes involved in heart development, thus providing a transcriptional correlate of the phenotypic observations. Among the rescued genes, we noted several that are implicated in epithelial-mesenchymal transition (EMT) and we selected *Snai2*-SLUG for further studies. At E9.5, SLUG identifies a mesodermal population in the PhAp that partially overlaps with other markers of the CPM, such as TBX1, ISL1, and Mef2c-AHF-Cre. This suggests that the SLUG+ cell population may include SHF cardiac progenitors and branchiomeric muscle progenitors of the cardiopharyngeal lineage. Importantly, the SLUG expression pattern changes in heterozygous and homozygous *Tbx1* mutants. This could be due to ectopic expression of the *Snai2* gene, or to mislocalization of SLUG+ cells. However, the second hypothesis, i.e. defective regionalization, is supported by the finding that the expression of other genes also follows similar pattern changes. In addition, the finding that NCCs, as identified by TFAP2A staining, are also mislocalized, supports the hypothesis that the *Tbx1* mutation is associated with anomalous regionalization of multiple cell lineages. It is unlikely that regionalization anomalies are due to morphogenetic defects because some of these anomalies, along with gene expression dysregulation, are also evident in the heterozygous mutants that do not show major morphogenetic defects.

The aggregation of SLUG+ cells, particularly evident in the 1st PA, and the segregation of these cells from the neural crest lineage, suggest that the mutation is altering mechanisms of cell sorting, a crucial process in embryonic morphogenesis. This problem may occur for a number of reasons. For example, the differential adhesion hypothesis (Steinberg, 1962) predicts that cells tend to group together if they have higher affinity with each other, compared with other neighboring cell populations. This possibility is supported by the finding that aggregated cells express higher levels of N-cadherin, a cell adhesion molecule, comared to the surrounding NCC-derived mesenchyme in the 1st PA of *Tbx1*^-/-^ emryo. Our observations in the 1st PA of early embryos indicate that in *Tbx1* mutants, incoming mesenchymal cells fail to mix with core mesodermal cells, consistently with a differential cell adhesion hypothesis. Intermingling of Myf5+ myogenic core cells and incoming TFAP2A+ NCCs in the 1st PA of WT embryos has previously been described (Grenier et al., 2009), although the mechanisms that govern this process are not yet established. We show here that TBX1 function is part of these mechanisms, although the effectors remain to be identified. The association of *Tbx1* heterozygosity and NCCs distribution has been noted previously in the posterior pharynx (Calmont et al., 2009). Cell -cell adhesion and/or cell-ECM interactions may interfere with NCCs migration delaying proper localization. Treatment with vB12 could also target NCCs and improve their migratory potential.

It is tempting to speculate that lineage regionalization abnormalities are part of the pathogenetic mechanism underlying the severe developmental defects of the PhAp associated with *Tbx1* mutation. Mislocalization, even transient, of different cell types may expose them to different signaling cues (or different concentrations thereof) causing further developmental defects downstream.

In this work, we used *Snai2* as a marker gene, but we did not address a potential role of *Snai2* in the observed phenotypes. SLUG is a transcriptional repressor that targets genes encoding adhesion molecules (such as e-cadherin) in epithelial cells, thus supporting their mobilization and mesenchymalization (Zhou et al., 2019). It is difficult to directly apply these concepts to the pharyngeal mesenchymal cells that we have studied. However, SLUG has also been associated with a number of different functions, including skeletal muscle differentiation (Tang et al., 2016) and with up-regulated N-cadherin in some contexts (review in (Loh et al., 2019)). Therefore, it would be of interest to determine in the future whether SLUG has a specific role in branchiomeric muscle differentiation or development, which is impaired in *Tbx1* mutants (Grifone et al., 2008; Kelly et al., 2004).

In summary, we show that vB12 treatment is sufficient to rescue in part several anomalies of the PhAp. We leveraged this activity to identify a set of genes, already known to be involved in heart development that may be part of or associated with the pathogenesis of TBX1-dependent phenotypes. Finally, one of these genes, encoding SLUG has been instrumental in the discovery of a novel phenotype of lineage regionalization defects.

## MATERIALS AND METHODS

### Mouse lines

In this work we used mouse lines previously described *Tbx1^lacZ^* (here referred to as *Tbx1*^+/-^) (Lindsay et al., 2001), *Tbx1^neo2^* (Zhang et al., 2006), *Tbx1^cre^* (Huynh et al., 2007), *Tbx1^flox^* (Xu et al., 2004), R26R^mTmG^ (Muzumdar et al., 2007), and Mef2c-AHF-Cre (Verzi et al., 2005). We have crossed *Tbx1^lacZ/+^* mice with *Tbx1^lacZ/+^* or *Tbx1^neo2^* to generate heterozygous, wild type, null or hypomorphic embryos. We have crossed *Tbx1^cre^* with R26R^mTmG^ mice to map distribution of *Tbx1*-expressing cells and their descendants. Administration of vB12 (cyanocobalamin Sigma-Aldrich Prod. Number V2876) was injected intraperitoneally (20mg/Kg). The impact of vB12 on great vessels and ventricular septation defects was scored at embryonic day (E) 15.5 and E18.5. Pregnant females were injected daily from E7.5 to E11.5. Developmental stages were assessed by considering the morning of vaginal plug as E0.5. Control mice were injected with the same volume of PBS.

Animal studies were carried out under the auspices of the animal protocol 257/2015-PR (licensed to the A.B. laboratory) reviewed, according to Italian regulations, by the Italian Istituto Superiore di Sanità and approved by the Italian Ministero della Salute.

### Mouse phenotyping

E15.5 and E18.5 embryos were examined under the stereomicroscope, fixed overnight in 4% paraformaldehyde (PFA) and E15.5 embryos embedded in paraffin, sectioned, and stained with eosin and hematoxylin. E18.5 hearts and great vessels were manually dissected and photographed under a stereomicroscope, and then embedded in paraffin, sectioned, and stained.

### Immunofluorescence

Embryos were fixed overnight in 4% PFA and embedded in wax. For immunofluorescence analysis embedded embryos were cutted in 7μm sections. Sections were deparaffinized in xilene, rehydrated, and after antigen unmasking with citrate buffer, sections were incubated overnight at room temperature with primary antibodies (in 0.5% milk, 10% fetal bovine serum, 1% bovine serum albumin in H2O). Each experiment was repeated at least three times.

We used the following primary antibodies: Anti-GFP (Abcam ab13970, diluted 1:1000), Anti-SLUG (Cell Signaling, #9585, diluted 1:100); Anti-TFAP2A (Hybridoma Bank, clone 3B5, diluted 1:300); Anti-ISL1, (Hybridoma Bank, clone 39.4D5, diluted 1:100); Anti-CRE (Millipore, 69050-3, diluted 1:1000); Anti-TBX1 (Abcam, Ab18530, diluted 1:100).

### RNA extraction and RT-PCR

Total RNA was isolated from E9.5 (22 somites) *Tbx1^+/-^* and *Tbx1^+/+^* embryos with TRIZOL (Invitrogen) and reverse-transcribed using the High Capacity cDNA reverse transcription kit (Applied Biosystem catalog. n. 4368814).

### RNA-seq gene expression data analysis

Processing and analysis RNA-Seq data: raw data for the high-throughput sequencing of cDNA were generated with Illumina platform for strand specific paired-end reads. These reads are 125bp long. In total 8 RNA-seq samples were sequenced. The three biological replicates of RNA-seq for *Tbx1*^+/-^ condition are indicated as Tbx1^+/-^ _rep1, Tbx1^+/-^ _rep2 and Tbx1^+/-^ _rep3, respectively. The two biological replicates of RNA-seq for heterozygous with B12 treatment are denoted *Tbx1^+/-^* (+vB12)_rep1 and *Tbx1^+/-^* (+vB12)_rep2 and the three biological replicates of RNA-seq for wild type condition are denoted Tbx1^+/+^ _rep1, Tbx1^+/+^ _rep2 and Tbx1^+/+^ _rep3, respectively. The quality control on raw reads was performed using FastQC (http://www.bioinformatics.babraham.ac.uk/projects/fastqc/).

Alignment of Sequence Reads: First, the reads were mapped to the mouse genome (mm9) using TopHat2 (version.2.0.7) (Trapnell et al., 2009), with the following options: -G annotation_file.gtf --transcriptome-index transcriptome. All other parameters were used with their default values. The annotation GTF (Gene transfer format) file, Mus_musculus.NCBIM37.67.gtf, was downloaded from http://www.ensembl.org.

Differential expressed genes (DEGs): Gene count matrix was obtained as output using featureCount function from Rsubread R package (version 0.5.4) on ‘exon’ feature type, considering reverse strand for paired end reads with the annotation GTF file. We selected the total counts on 37620 genes for differential expression analysis.

The raw counts were filtered applying Proportion test or a total of 14488 genes RNA-SeqGUI R package (Russo and Angelini, 2014). Principal component analysis (PCA) was performed to separate biological conditions. PCA results showed that samples clustered for different library preparations and different times and therefore raw data had to be corrected for batch effects. Firstly, we performed a complex design, considering that exist a not identified batch. We removed batch effects using ARSyNseq function from filtered gene count matrix and considering rpkm normalization approach. Then, we evaluated differential expression between pair-conditions using the non-parametric NOISeqBIO function (Tarazona et al., 2015) after applying upper quartile as normalization method. A posterior probability (PP) greater or equal to 0.95 was used to determine DEGs.

The list of DE genes with absolute value of fold-change greater or equal 1.2 were considered for pathway analysis. In addition, we performed the pathway analysis using the g:Profiler tool (Raudvere et al., 2019), setting the organism to *mus musculus,* choosing as custom background the list of 14488 expressed genes in our system, setting the significance threshold for the multiplicity correction “fdr” (i.e, Benjamini and Hochberg FDR) with the user threshold 0.05. We limit the sources to GO, KEGG, and Human phenotype ontology databases to evaluate functional enrichment.

## ACKNOWLEDGEMENTS

We are grateful to all members of our laboratories and colleagues of the Leducq network for helpful discussion. We also thank Rosa Ferrentino for expert laboratory assistance and Elizabeth Illingworth for critical reading of the manuscript. We thanks Salvatore Arbucci for his technical support for for the acquisition of images under the confocal microscope and IGB microscopy facility. We thanks Lucia Mele for her technical support in mouse treatments and IGB mouse facility.

This work was funded by the Fondation Leducq Transatlantic Network of Excellence 15CVD01 (to AB and RGK), a grant from the Italian Ministry of University and Research PRIN 20179[2P9] (to AB).

## AUTHOR CONTRIBUTIONS

AB: Conceptualization, Writing - Original Draft, Supervision, Funding Acquisition; AN: Investigation, Formal analysis, Writing - Review & Editing. AR, ED, and MB: Investigation; CA: Formal analysis, Supervision, Writing - Review & Editing; GL: Conceptualization, Investigation, Formal analysis, Supervision, Writing - Review & Editing; MF: Formal analysis; RGK: Funding Acquisition, Writing - Review & Editing.

## REFERENCES

Adachi N, Bilio M, Baldini A, Kelly RG. 2020. Cardiopharyngeal mesoderm origins of musculoskeletal and connective tissues in the mammalian pharynx. Development 147:10.1242/dev.185256. doi:10.1242/dev.185256

Alimperti S, Andreadis ST. 2015. CDH2 and CDH11 act as regulators of stem cell fate decisions. Stem Cell Res 14:270–282. doi:10.1016/j.scr.2015.02.002

Calmont A, Ivins S, Van Bueren KL, Papangeli I, Kyriakopoulou V, Andrews WD, Martin JF, Moon AM, Illingworth EA, Basson MA, Scambler PJ. 2009. Tbx1 controls cardiac neural crest cell migration during arch artery development by regulating Gbx2 expression in the pharyngeal ectoderm. Development 136:3173–83.

Dastjerdi A, Robson L, Walker R, Hadley J, Zhang Z, Rodriguez-Niedenfuhr M, Ataliotis P, Baldini A, Scambler P, Francis-West P. 2007. Tbx1 regulation of myogenic differentiation in the limb and cranial mesoderm. Dev Dyn 236:353–63.

Fulcoli FG, Franzese M, Liu X, Zhang Z, Angelini C, Baldini A. 2016. Rebalancing gene haploinsufficiency in vivo by targeting chromatin. Nat Commun 7:11688. doi:10.1038/ncomms11688

Graham A. 2001. The development and evolution of the pharyngeal arches. J Anat 199:133–41.

Grenier J, Teillet M-A, Grifone R, Kelly RG, Duprez D. 2009. Relationship between neural crest cells and cranial mesoderm during head muscle development. PloS One 4:e4381. doi:10.1371/journal.pone.0004381

Grifone R, Jarry T, Dandonneau M, Grenier J, Duprez D, Kelly RG. 2008. Properties of branchiomeric and somite-derived muscle development in Tbx1 mutant embryos. Dev Dyn Off Publ Am Assoc Anat 237:3071–3078. doi:10.1002/dvdy.21718

Haddad RA, Clines GA, Wyckoff JA. 2019. A case report of T-box 1 mutation causing phenotypic features of chromosome 22q11.2 deletion syndrome. Clin Diabetes Endocrinol 5:13. doi:10.1186/s40842-019-0087-6

Huang GY, Cooper ES, Waldo K, Kirby ML, Gilula NB, Lo CW. 1998. Gap junction-mediated cell-cell communication modulates mouse neural crest migration. J Cell Biol 143:1725–34.

Huynh T, Chen L, Terrell P, Baldini A. 2007. A fate map of Tbx1 expressing cells reveals heterogeneity in the second cardiac field. Genesis 45:470–5.

Ivins S, Lammerts van Beuren K, Roberts C, James C, Lindsay E, Baldini A, Ataliotis P, Scambler PJ. 2005. Microarray analysis detects differentially expressed genes in the pharyngeal region of mice lacking Tbx1. Dev Biol 285:554–569. doi:10.1016/j.ydbio.2005.06.026

Jerome LA, Papaioannou VE. 2001. DiGeorge syndrome phenotype in mice mutant for the T-box gene, Tbx1. Nat Genet 27:286–291.

Kelly RG, Jerome-Majewska LA, Papaioannou VE. 2004. The del22q11.2 candidate gene Tbx1 regulates branchiomeric myogenesis. Hum Mol Genet 13:2829–2840. doi:10.1093/hmg/ddh304

Kodo K, Shibata S, Miyagawa-Tomita S, Ong S-G, Takahashi H, Kume T, Okano H, Matsuoka R, Yamagishi H. 2017. Regulation of Sema3c and the Interaction between Cardiac Neural Crest and Second Heart Field during Outflow Tract Development. Sci Rep 7:6771. doi:10.1038/s41598-017-06964-9

Lania G, Bresciani A, Bisbocci M, Francone A, Colonna V, Altamura S, Baldini A. 2016. Vitamin B12 ameliorates the phenotype of a mouse model of DiGeorge syndrome. Hum Mol Genet 25:4369–4375. doi:10.1093/hmg/ddw267

Liao J, Aggarwal VS, Nowotschin S, Bondarev A, Lipner S, Morrow BE. 2008. Identification of downstream genetic pathways of Tbx1 in the second heart field. Dev Biol 316:524–37.

Liao J, Kochilas L, Nowotschin S, Arnold JS, Aggarwal VS, Epstein JA, Brown MC, Adams J, Morrow BE. 2004. Full spectrum of malformations in velo-cardio-facial syndrome/DiGeorge syndrome mouse models by altering Tbx1 dosage. Hum Mol Genet 13:1577–85.

Lindsay EA, Vitelli F, Su H, Morishima M, Huynh T, Pramparo T, Jurecic V, Ogunrinu G, Sutherland HF, Scambler PJ, Bradley A, Baldini A. 2001. Tbx1 haploinsufficieny in the DiGeorge syndrome region causes aortic arch defects in mice. Nature 410:97–101.

Loh C-Y, Chai JY, Tang TF, Wong WF, Sethi G, Shanmugam MK, Chong PP, Looi CY. 2019. The E-Cadherin and N-Cadherin Switch in Epithelial-to-Mesenchymal Transition: Signaling, Therapeutic Implications, and Challenges. Cells 8:E1118. doi:10.3390/cells8101118

Mao A, Zhang M, Li L, Liu J, Ning G, Cao Y, Wang Q. 2021. Pharyngeal pouches provide a niche microenvironment for arch artery progenitor specification. Dev Camb Engl 148:dev192658. doi:10.1242/dev.192658

Merscher S, Funke B, Epstein JA, Heyer J, Puech A, Min Lu MM, Xavier RJ, Demay MB, Russell RG, Factor S, Tokooya K, St. Jore B, Lopez M, Pandita RK, Lia M, Carrion D, Xu H, Schorle H, Kobler JB, Scambler PJ, Wynshaw-Boris A, Skoultchi AI, Morrow BE, Kucherlapati R. 2001. TBX1 Is Responsible for Cardiovascular Defects in Velo-Cardio-Facial/DiGeorge Syndrome. Cell 104:619–629.

Muzumdar MD, Tasic B, Miyamichi K, Li L, Luo L. 2007. A global double-fluorescent Cre reporter mouse. Genes N Y N 2000 45:593–605. doi:10.1002/dvg.20335

Pane LS, Zhang Z, Ferrentino R, Huynh T, Cutillo L, Baldini A. 2012. Tbx1 is a negative modulator of Mef2c. Hum Mol Genet 21:2485–96. doi:10.1093/hmg/dds063

Paylor R, Glaser B, Mupo A, Ataliotis P, Spencer C, Sobotka A, Sparks C, Choi CH, Oghalai J, Curran S, Murphy KC, Monks S, Williams N, O’Donovan MC, Owen MJ, Scambler PJ, Lindsay E. 2006. Tbx1 haploinsufficiency is linked to behavioral disorders in mice and humans: implications for 22q11 deletion syndrome. Proc Natl Acad Sci U A 103:7729–34.

Raudvere U, Kolberg L, Kuzmin I, Arak T, Adler P, Peterson H, Vilo J. 2019. g:Profiler: a web server for functional enrichment analysis and conversions of gene lists (2019 update). Nucleic Acids Res 47:W191–W198. doi:10.1093/nar/gkz369

Russo F, Angelini C. 2014. RNASeqGUI: a GUI for analysing RNA-Seq data. Bioinforma Oxf Engl 30:2514–2516. doi:10.1093/bioinformatics/btu308

Sato A, Scholl AM, Kuhn EN, Kuhn EB, Stadt HA, Decker JR, Pegram K, Hutson MR, Kirby ML. 2011. FGF8 signaling is chemotactic for cardiac neural crest cells. Dev Biol 354:18–30. doi:10.1016/j.ydbio.2011.03.010

Shone V, Graham A. 2014. Endodermal/ectodermal interfaces during pharyngeal segmentation in vertebrates. JAnat 225:479–491. doi:10.1111/joa.12234

Steinberg MS. 1962. On the mechanism of tissue reconstruction by dissociated cells. I. Population kinetics, differential adhesiveness. and the absence of directed migration. Proc Natl Acad Sci U S A 48:1577–1582. doi:10.1073/pnas.48.9.1577

Swedlund B, Lescroart F. 2020. Cardiopharyngeal Progenitor Specification: Multiple Roads to the Heart and Head Muscles. Cold Spring Harb Perspect Biol 12:a036731. doi:10.1101/cshperspect.a036731

Tang Y, Feinberg T, Keller ET, Li X-Y, Weiss SJ. 2016. Snail/Slug binding interactions with YAP/TAZ control skeletal stem cell self-renewal and differentiation. Nat Cell Biol 18:917–929. doi:10.1038/ncb3394

Tarazona S, Furió-Tarí P, Turrà D, Pietro AD, Nueda MJ, Ferrer A, Conesa A. 2015. Data quality aware analysis of differential expression in RNA-seq with NOISeq R/Bioc package. Nucleic Acids Res 43:e140. doi:10.1093/nar/gkv711

Trapnell C, Pachter L, Salzberg SL. 2009. TopHat: discovering splice junctions with RNA-Seq. Bioinforma OxfEngl 25:1105–1111. doi:10.1093/bioinformatics/btp120

Verzi MP, McCulley DJ, De Val S, Dodou E, Black BL. 2005. The right ventricle, outflow tract, and ventricular septum comprise a restricted expression domain within the secondary/anterior heart field. Dev Biol 287:134–45.

Wang W, Niu X, Stuart T, Jullian E, Mauck WM, Kelly RG, Satija R, Christiaen L. 2019. A single-cell transcriptional roadmap for cardiopharyngeal fate diversification. Nat Cell Biol 21:674–686. doi:10.1038/s41556-019-0336-z

Warkala M, Chen D, Ramirez A, Jubran A, Schonning MJ, Wang X, Zhao H, Astrof S. 2020. Cell - ECM Interactions Play Multiple Essential Roles in Aortic Arch Development. Circ Res. doi:10.1161/CIRCRESAHA.120.318200

Xu H, Morishima M, Wylie JN, Schwartz RJ, Bruneau BG, Lindsay EA, Baldini A. 2004. Tbx1 has a dual role in the morphogenesis of the cardiac outflow tract. Development 131:3217–27.

Xu Y-J, Chen S, Zhang J, Fang S-H, Guo Q-Q, Wang J, Fu Q-H, Li F, Xu R, Sun K. 2014. Novel TBX1 loss-of-function mutation causes isolated conotruncal heart defects in Chinese patients without 22q11.2 deletion. BMCMed Genet 15:78. doi:10.1186/1471-2350-15-78

Yagi H, Furutani Y, Hamada H, Sasaki T, Asakawa S, Minoshima S, Ichida F, Joo K, Kimura M, Imamura S, Kamatani N, Momma K, Takao A, Nakazawa M, Shimizu N, Matsuoka R. 2003. Role of TBX1 in human del22q11.2 syndrome. Lancet 362:1366–1373.

Zhang Z, Baldini A. 2008. In vivo response to high-resolution variation of Tbx1 mRNA dosage. Hum Mol Genet 17:150–7.

Zhang Z, Huynh T, Baldini A. 2006. Mesodermal expression of Tbx1 is necessary and sufficient for pharyngeal arch and cardiac outflow tract development. Development 133:3587–3595.

Zhou W, Gross KM, Kuperwasser C. 2019. Molecular regulation of Snai2 in development and disease. J Cell Sci 132. doi:10.1242/jcs.235127

Zweier C, Sticht H, Aydin-Yaylagul I, Campbell CE, Rauch A. 2007. Human TBX1 missense mutations cause gain of function resulting in the same phenotype as 22q11.2 deletions. Am J Hum Genet 80:510–7.

