## Supplementary figures and Tab1 for "A phenotypic rescue approach identifies lineage regionalization defects in a mouse model of DiGeorge syndrome"

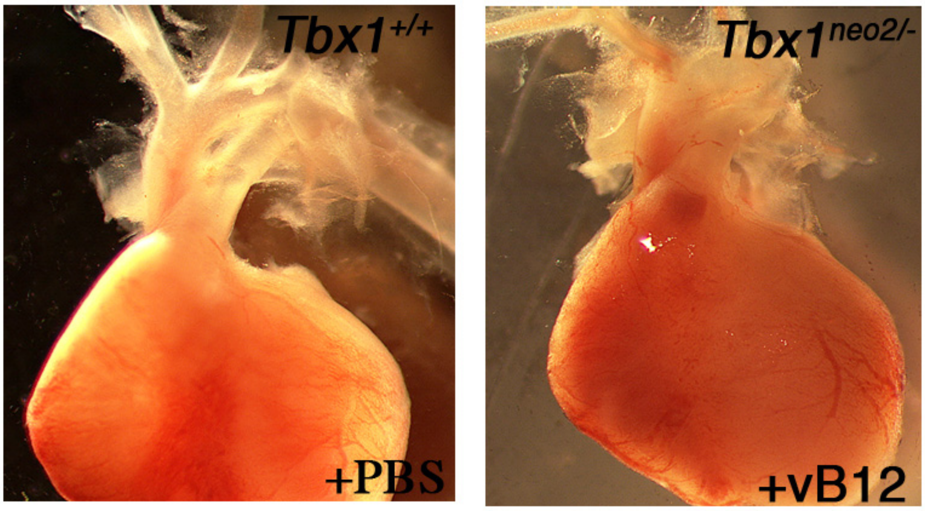


**Supplementary Figure 1** *Whole mount photographs of E18.5 hearts isolated from* $Tbx1^{+/+}$ *and* $Tbx1^{neo2/-}$ *treated with vB12*. The two hearts are apparently indistinguishable. Atria have been removed to show the proximal great arteries.

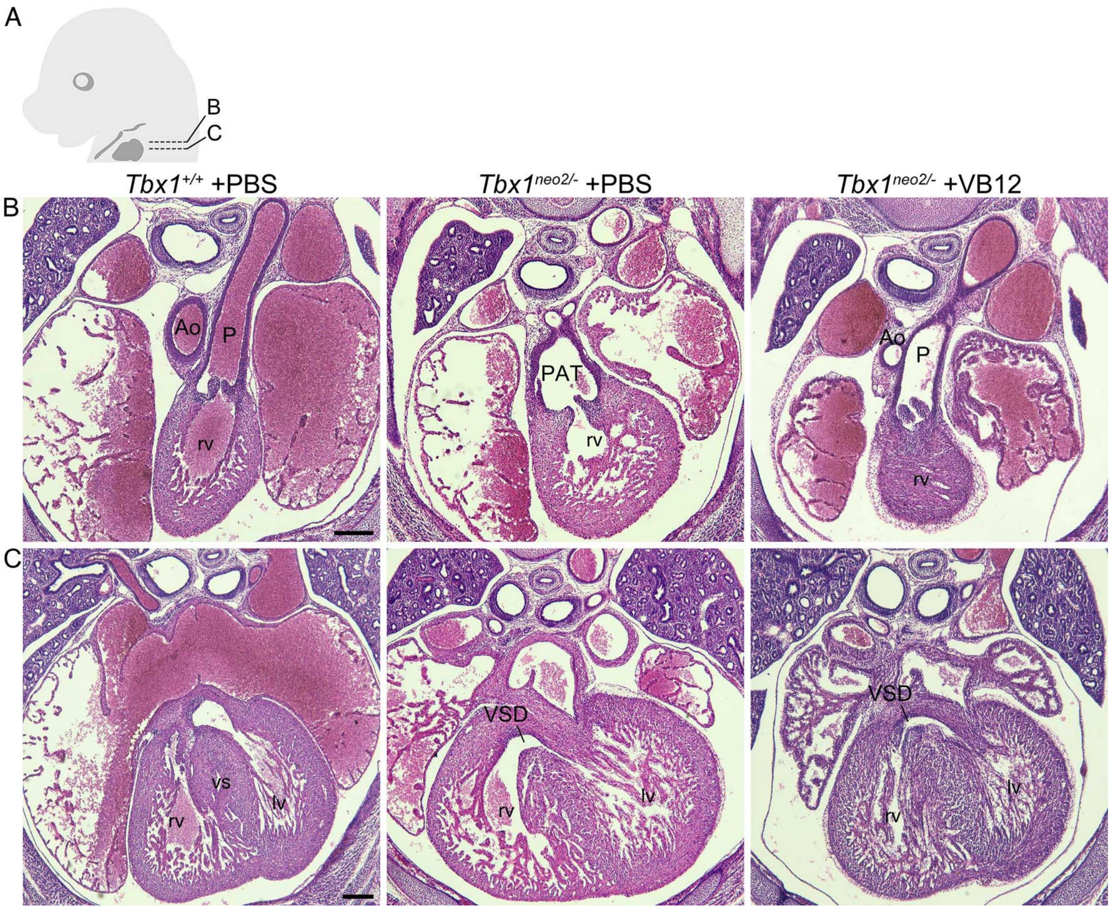


**Supplementary Figure 2** *Vitamin B12 improves cardiac outflow tract defects observed in Tbx1$^{neo2/-}$ mouse embryos at E15.5.* A) A diagram showing the section levels. B-C) Transverse histological sections of the heart from *Tbx1$^{+/+}$* and *Tbx1$^{neo2/-}$* embryos at E15.5, treated with PBS or vitamin B12. B) Outlet and C) ventricular septal levels. After vB12 treatment, *Tbx1$^{neo2/-}$* embryos show separated aorta (Ao) and pulmonary trunk (P), but have overriding aorta (OAo).

lv, left ventricle; PTA, Persistent truncus arteriosus; rv, right ventricle; vs, ventricular septum; VSD, ventricular septal defect.

Scale bar: 200 μm.

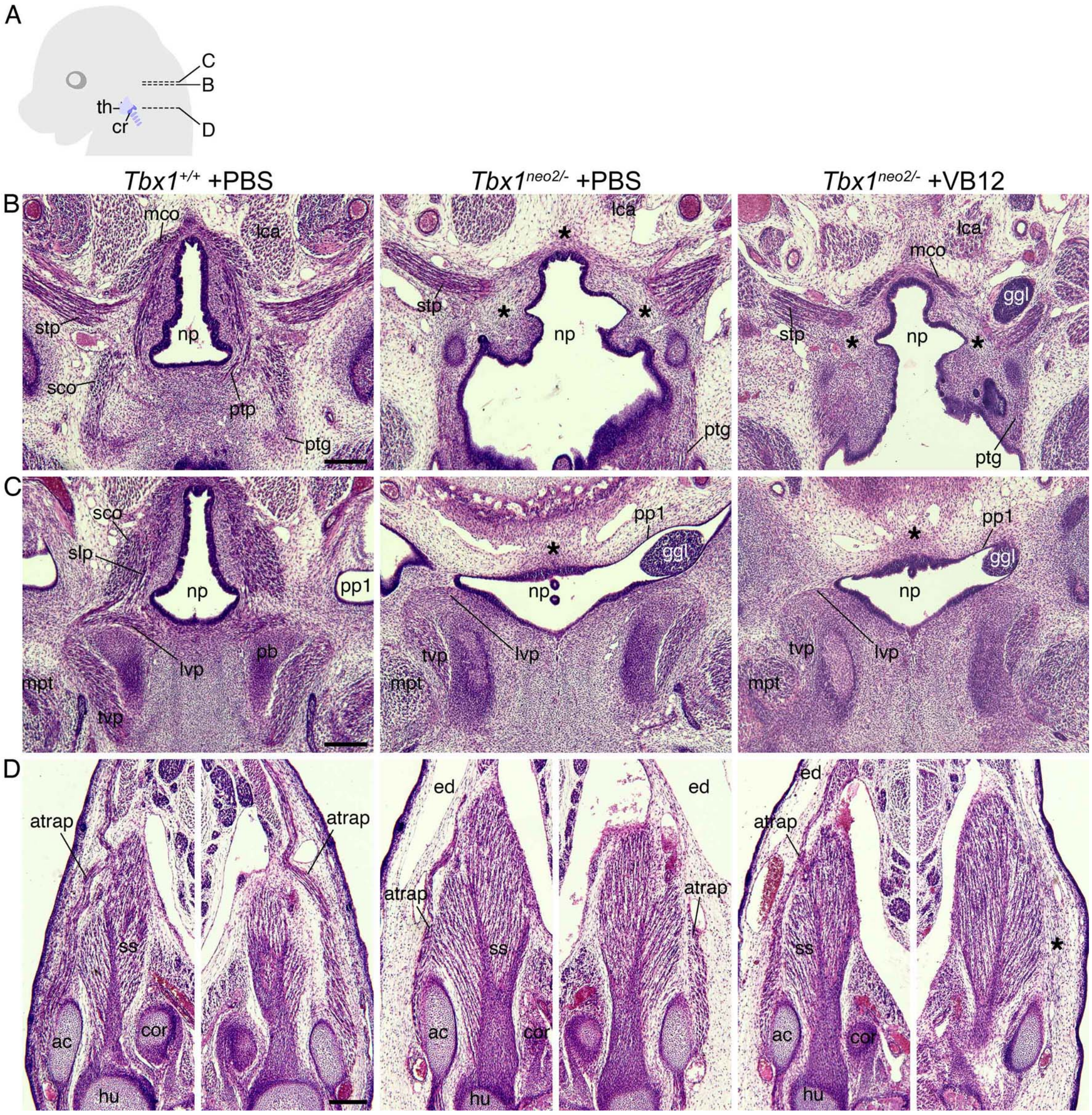


**Supplementary Figure 3** *Vitamin B12 treatment has little effect on the development of branchiomeric muscles derived from 3-6 pharyngeal arches in Tbx1$^{neo2/-}$ embryos.* A) A diagram showing the section levels. B-D) Transverse histological sections of *Tbx1*$^{+/+}$ and *Tbx1*$^{neo2/-}$ embryos at E15.5, treated with PBS or vitamin B12. B-C) Nasopharynx and D) shoulder levels. The asterisks indicate missing muscles. Branchiomeric muscles defective in *Tbx1*$^{neo2/-}$ embryos are not clearly rescued after VB12 treatment. Some muscles exhibit normal development even in the low expression level of *Tbx1*.

ac, acromion; ary, arytenoid cartilage; atrap, acromiotrapezius muscle; cor, coracoid process; ed, edema; ggl, geniculate ganglion-like tissue; hu, humerus; lca, longs capitis muscle; lvp, levator veli palatini muscle; mco, middle constrictor muscle; mpt, medial pterygoid muscle; np, nasopharynx; pb, palatine bone; pp1, first pharyngeal pouch; ptg, palatoglossus muscle; ptp, palatopharyngeus muscle; sco, superior constrictor muscle; slp, salpingopharyngeus muscle; ss, supraspinatus muscle; stp, stylopharyngeus muscle; tvp, tensor veli palati muscle.

Scale bar: 200 μm.

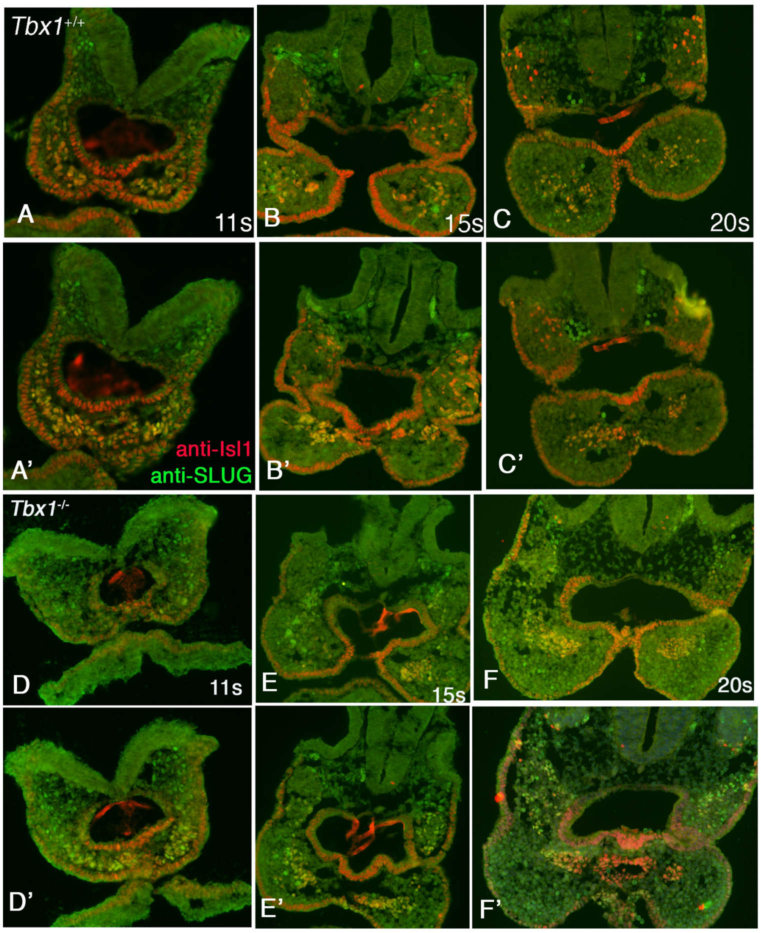


**Supplementary Figure 4** *SLUG+ and ISL1+ cells of the 1st pharyngeal arch of Tbx1$^{-/-}$ embryos fail to intermingle with the arch mesenchyme.*

Immunofluorescence analysis of SLUG and ISL1 at 11, 15 and 22 somites stages of *Tbx1*$^{+/+}$ (A-C) and *Tbx1*$^{-/-}$ (D-F) embryos. Representative images of two consecutive sections of most cranial part of the embryos, at the level of the 1st and 2nd pharyngeal arches.

Scale bar: 100 μm

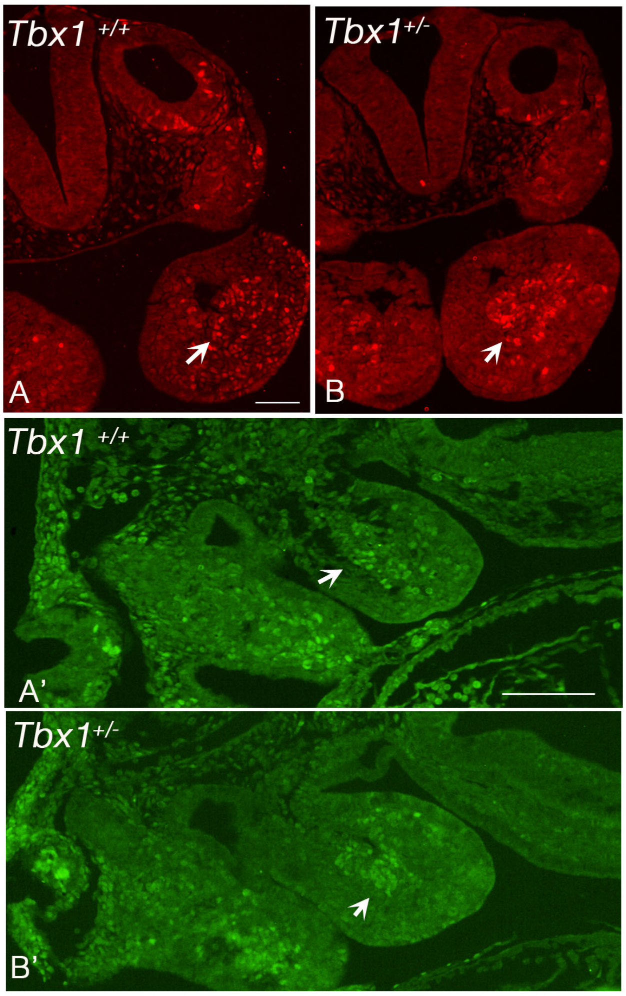


**Supplementary Figure 5** *Increased mesodermal cell condensation in the 1st pharyngeal arch of Tbx1 heterozygous embryos at E9.5.*

Immunofluorescence of transverse (top panels, in red) and sagittal lateral (bottom panels in green) using anti-SLUG antibodies. Arrows point to a group of SLUG+ cells in the 1st pharyngeal arch. Note the higher condensation of SLUG+ cells in the heterozygous mutant.

Scale bar: 100 μm

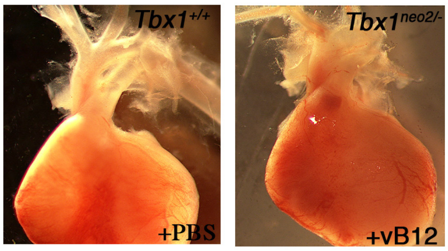


**Supplementary Figure 1** *Whole mount photographs of E18.5 hearts isolated from $Tbx1^{+/+}$ and $Tbx1^{neo2/-}$ treated with vB12*. The two hearts are apparently indistinguishable. Atria have been removed to show the proximal great arteries.

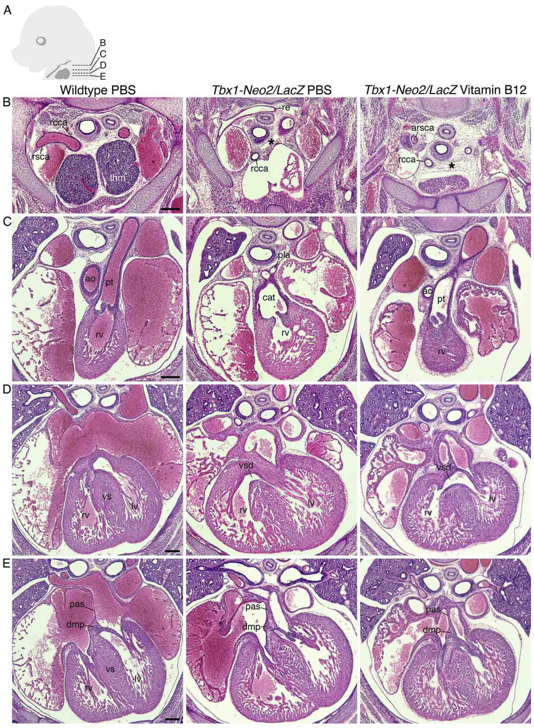


**Supplementary Figure 2** *Vitamin B12 improves cardiac outflow tract defects observed in $Tbx1^{neo2/-}$ mouse embryos at E15.5.* A) A diagram showing the section levels. B-C) Transverse histological sections of the heart from $Tbx1^{+/+}$ and $Tbx1^{neo2/-}$ embryos at E15.5, treated with PBS or vitamin B12. B) Outlet and C) ventricular septal levels. After vB12 treatment, $Tbx1^{neo2/-}$ embryos show separated aorta (Ao) and pulmonary trunk (P), but have overriding aorta (OAo).

lv, left ventricle; PTA, Persistent truncus arteriosus; rv, right ventricle; vs, ventricular septum; VSD, ventricular septal defect.

Scale bar: 200 µm.

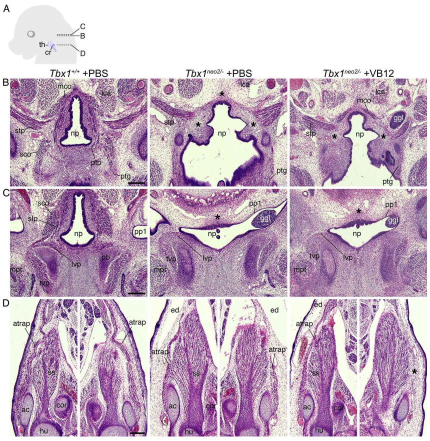


**Supplementary Figure 3** *Vitamin B12 treatment has little effect on the development of branchiomeric muscles derived from 3-6 pharyngeal arches in Tbx1$^{neo2/-}$ embryos.* A) A diagram showing the section levels. B-D) Transverse histological sections of *Tbx1*$^{+/+}$ and *Tbx1*$^{neo2/-}$ embryos at E15.5, treated with PBS or vitamin B12. B-C) Nasopharynx and D) shoulder levels. The asterisks indicate missing muscles. Branchiomeric muscles defective in *Tbx1*$^{neo2/-}$ embryos are not clearly rescued after VB12 treatment. Some muscles exhibit normal development even in the low expression level of *Tbx1*.

ac, acromion; ary, arytenoid cartilage; atrap, acromiotrapezius muscle; cor, coracoid process; ed, edema; ggl, geniculate ganglion-like tissue; hu, humerus; lca, longs capitis muscle; lvp, levator veli palatini muscle; mco, middle constrictor muscle; mpt, medial pterygoid muscle; np, nasopharynx; pb, palatine bone; pp1, first pharyngeal pouch; ptg, palatoglossus muscle; ptp, palatopharyngeus muscle; sco, superior constrictor muscle; slp, salpingopharyngeus muscle; ss, supraspinatus muscle; stp, stylopharyngeus muscle; tvp, tensor veli palati muscle.

Scale bar: 200 µm.

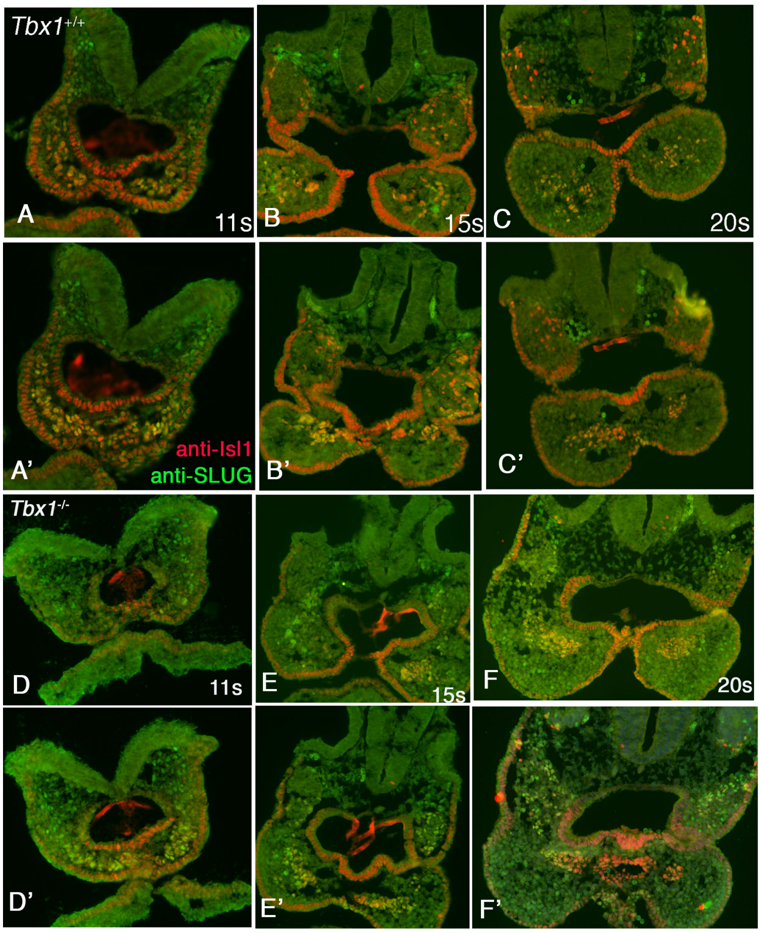


**Supplementary Figure 4** *SLUG+ and ISL1+ cells of the 1st pharyngeal arch of Tbx1$^{-/-}$ embryos fail to intermingle with the arch mesenchyme.*

Immunofluorescence analysis of SLUG and ISL1 at 11, 15 and 22 somites stages of *Tbx1*$^{+/+}$ (A-C) and *Tbx1*$^{-/-}$ (D-F) embryos. Representative images of two consecutive sections of most cranial part of the embryos, at the level of the 1st and 2nd pharyngeal arches.

Scale bar: 100 μm

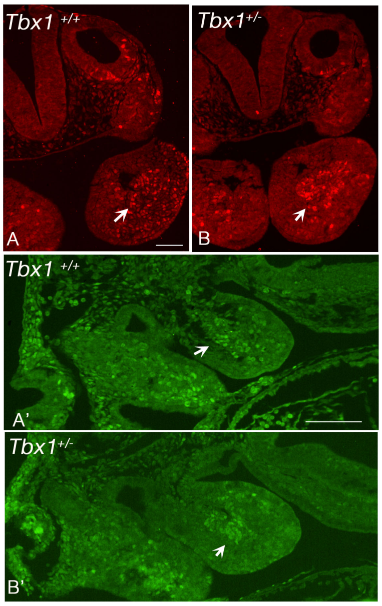


**Supplementary Figure 5** *Increased mesodermal cell condensation in the 1st pharyngeal arch of Tbx1 heterozygous embryos at E9.5.*

Immunofluorescence of transverse (top panels, in red) and sagittal lateral (bottom panels in green) using anti-SLUG antibodies. Arrows point to a group of SLUG+ cells in the 1st pharyngeal arch. Note the higher condensation of SLUG+ cells in the heterozygous mutant.

Scale bar: 100 μm

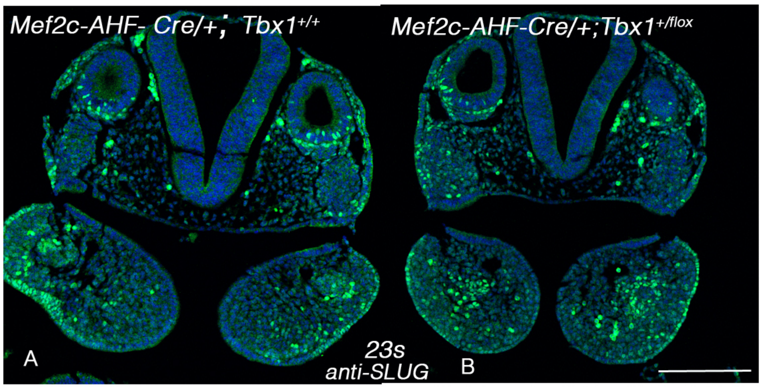


**Supplementary Figure 6** *The condensation phenotype is detectable in conditional mutants.*Anti-SLUG immunofluorescence of transverse sections through the 1st pharyngeal arch of E9.5 embryos. The condensation of SLUG+ cells in the core of the arch is also evident in the Mef2c-AHF-Cre;*Tbx1*$^{flox/+}$ mutant. Scale bar: 200 μm

### Supplementary Table 1

*Banchiomeric muscle phenotype*

| Branchiomeric muscles | | n | Treatment | Normal | Unilateral defect | Bilateral defect |
|---|---|---|---|---|---|---|
| ***Tbx1$^{neo2/-}$*** **1st PA muscle** | Ma, Mh, Pt, Te, Tt, Tvp | 5 | PBS | 5 | 0 | 0 |
| | Ma, Mh, Pt, Te, Tt, Tvp | 3 | VB12 | 3 | 0 | 0 |
| | Ad | 5 | PBS | 0 | 2 | 3 |
| | Ad | 3 | VB12 | 1 | 1 | 1 |
| ***Tbx1$^{neo2/-}$*** **2nd PA muscle** | Fe | 5 | PBS | 5 | 0 | 0 |
| | Fe | 3 | VB12 | 3 | 0 | 0 |
| | Stm | 5 | PBS | 1 | 1 | 3 |
| | Stm | 3 | VB12 | 1 | 2 | 0 |
| | Pd | 5 | PBS | 1 | 3 | 1 |
| | Pd | 3 | VB12 | 1 | 2 | 0 |
| | Sty | 5 | PBS | 0 | 0 | 5 |
| | Sty | 3 | VB12 | 0 | 0 | 3 |
| ***Tbx1$^{neo2/-}$*** **3rd PA muscle** | Stp | 5 | PBS | 5 | 0 | 0 |
| | Stp | 3 | VB12 | 3 | 0 | 0 |
| ***Tbx1$^{neo2/-}$*** **4-6th PA muscle** | Crth, Ic, Lvp, Ptg | 5 | PBS | 5 | 0 | 0 |
| | Crth, Ic, Lvp, Ptg | 3 | VB12 | 3 | 0 | 0 |
| | Thar | 5 | PBS | 4 | 0 | 1 |
| | Thar | 3 | VB12 | 3 | 0 | 0 |
| | Otary | 5 | PBS | 1 | 1 | 3 |
| | Otary | 3 | VB12 | 2 | 0 | 1 |
| | Lcary, Mc, Pcary | 5 | PBS | 0 | 1 | 4 |
| | Lcary, Mc, Pcary | 3 | VB12 | 0 | 0 | 3 |
| | Scm | 5 | PBS | 0 | 2 | 3 |
| | Scm | 3 | VB12 | 0 | 0 | 3 |
| | Ptp, Slp, Sc, Trap, Vm | 5 | PBS | 0 | 0 | 5 |
| | Ptp, Slp, Sc, Trap, Vm | 3 | VB12 | 0 | 0 | 3 |

Ad, anterior digastric muscle; Crth, cricothyroid muscle; Fe, facial expression muscle; Ic, inferior constrictor muscle; Lcary, lateral cricoarytenoid muscle; Lvp, levator veli palatini muscle; Ma, masseter muscle; Mc, middle constrictor muscle; Mh, mylohyoid muscle; Otary, oblique and transverse arytenoid muscle; Pcary, posterior cricoarytenoid muscle; Pd, posterior digastric muscle; Pt, pterygoid muscle; Ptg, palatoglossus muscle; Ptp, palatopharyngeus muscle; Sc, superior constrictor muscle; Scm, sternocleidomastoid muscle; Slp, salpingopharyngeus muscle; Stm, stapedius muscle; Stp, stylopharyngeus muscle; Sty, stylohyoid muscle; Te, temporal muscle; Thary, thyroarytenoid muscle; trap, trapezius muscle; Tt, tensor tympani muscle; Tvp, tensor veli palati muscle; Vm, vocal muscle.
